# Interpreting Short Tandem Repeat Variations with Genomic Foundation Models in Olive

**DOI:** 10.64898/2026.10.21.760403

**Authors:** Feifei Xia, Mohammad Hossein Pakdel, Mehdi Hosseini Mazinani, AliAkbar Asadi, Yanfei Zeng, Jianguo Zhang, Vahid Shariati, Maria Anisimova

## Abstract

Short tandem repeats (STRs) are highly mutable, multiallelic variants with distinctive mutation dynamics and have been associated with phenotypic variation across diverse species. However, genome-wide STR variation remains poorly characterized in olive, and the large number of naturally occurring alleles creates a challenge for variant interpretation. Genomic language models offer a potential way to assess how STR length changes alter learned sequence representations, but their responses to natural multiallelic STR series have not been systematically evaluated. Here, we identified 117,119 polymorphic STRs across 212 Iranian and Mediterranean olive accessions. STRs recovered the major SNP-defined ancestry axes while resolving additional geographically structured subcomponents. We then quantified STR-induced embedding perturbation using three plant genomic language models: AgroNT, PlantCAD2, and GPN. Across all three models, embedding distance generally increased with repeat-length difference from the reference allele, and 7,417 loci were embedding-sensitive in all three models. These signals were robust to fixed-length input construction and exceeded those of matched nonrepetitive indels at most tested loci. Integration with oil-composition associations prioritized four exploratory loci supported by all three models. Overall, we provide a genome-wide olive STR resource and show that plant genomic language models capture reproducible responses to natural STR length variation. Combining cross-model embedding evidence with phenotype association provides a potential framework for prioritizing candidate STRs for further investigation.

## Background

The olive tree (*Olea europaea* L.) is among the oldest perennial crops and remains central to agriculture throughout the Mediterranean basin and adjoining regions. Its present-day diversity reflects a complex history of lineage differentiation, domestication, recurrent admixture, and gene flow between cultivated and wild populations, rather than a single domestication trajectory^1–3^. Genome-wide studies have revealed substantial diversity within Mediterranean germplasm, while accessions from Iran and the eastern Fertile Crescent capture differentiation extending beyond the traditionally emphasized western and central Mediterranean gene pools^4–6^. This combination of deep population structure, recent genetic exchange, and extensive standing variation makes olive an informative system for studying crop evolution. It also complicates genotype–phenotype analyses, because alleles associated with ancestry, domestication history, and agronomic traits may be correlated across heterogeneous germplasm.

Reference genomes, population resequencing, quantitative trait locus mapping, and transcriptomic studies have substantially expanded the genomic resources available for olive improvement. Studies of oil composition have identified genomic regions and expression programs associated with variation in oleic, linoleic, and related fatty acids, including key roles for fatty-acid desaturase pathways^7–10^. Nevertheless, converting statistical associations and molecular observations into robust, breeding-relevant hypotheses remains a major challenge in crop genomics^11^. Population structure, linkage disequilibrium, incomplete functional annotation, and the long generation time of perennial crops all widen the gap between variant discovery and experimental validation. Moreover, most genome-wide studies focus primarily on single-nucleotide polymorphisms (SNPs) and short indels. Although these variants provide an essential view of olive diversity, other forms of variation with distinct mutation processes and allelic architectures may contribute complementary information about population history and quantitative traits.

Short tandem repeats (STRs) are a particularly relevant but comparatively undercharacterized component of this variation. STRs consist of tandemly repeated short sequence motifs and mutate predominantly through gains or losses of repeat units, generating multiallelic series of repeat lengths^12,13^. Their mutation rates and stepwise allele structure differ markedly from those of SNPs, creating a dynamic source of standing genetic variation. Repeat-length changes can also alter local sequence properties by modifying transcription-factor binding, gene expression, RNA-associated regulation, or other regulatory and transcribed features^14–16^. In plant, evidence from natural *Arabidopsis thaliana* populations further links STR length variation to gene expression and suggests that STR mutations can explain a measurable component of complex-trait heritability^17,18^. These findings make STRs plausible contributors to regulatory and phenotypic variation in crops, although genomic position or length variability alone does not establish a functional effect at an individual locus.

STRs have long been used as markers of olive diversity, but conventional microsatellite panels typically assay only a small number of selected loci^19,20^. Genome-wide genotyping instead makes it possible to analyze thousands of loci while retaining their multiallelic structure. Because STRs and SNPs differ in mutation rate and allelic state space, they need not capture identical aspects of population history. Genome-wide STR analyses in other systems have demonstrated their value for genetic-diversity assessment, and scalable latent-factor approaches can accommodate the structure of large multiallelic genotype matrices^21,22^. Recent work further indicates that genome-wide STRs can recover broad SNP-defined ancestry while subdividing some ancestry components^23^. A genome-wide olive STR panel could therefore complement established SNP resources by improving the resolution of fine-scale population structure and enabling association analyses of quantitative agronomic traits.

Interpreting the large number of alleles in such a panel presents an additional challenge, particularly for STRs outside well-annotated coding regions. Genomic foundation models learn representations of sequence context from large collections of unlabeled genomes and have shown promise for functional annotation and variant-effect prediction^24,25^. Plant-focused models, including AgroNT, GPN, and PlantCAD2, extend this framework across diverse plant genomes and sequence scales^26–28^. Their learned representations may therefore provide a complementary source of evidence for prioritizing repeat variants and linking sequence variation to crop-relevant phenotypes^29^. However, genomic foundation models have generally been evaluated using nucleotide substitutions, conventional indels, or downstream annotation tasks. To our knowledge, their responses to naturally occurring multiallelic STR length series have not been systematically examined. STRs provide a distinct test because changes in repeat copy number simultaneously alter motif dosage, local sequence length, and the spacing between flanking elements. Any resulting signal may also depend on model architecture, representation layer, input design, and distance metric. Embedding perturbation therefore offers a promising framework for identifying STR allelic changes that are consistently recognized by genomic foundation models and for prioritizing loci for downstream functional investigation.

Here, we analyzed whole-genome sequencing data from 212 Iranian and Mediterranean olive accessions to establish a genome-wide catalogue of polymorphic STRs and characterize their motif composition, genomic distribution, and repeat-length variation. We compared STR- and SNP-derived population structure to determine whether multiallelic repeat variation captured ancestry patterns beyond those resolved by SNPs, and we tested STR associations with oil fatty-acid composition. We then represented naturally observed STR alleles using three independently trained plant genomic foundation models, AgroNT, PlantCAD2, and GPN, and quantified reference-to-alternative changes in embedding space. Cross-model comparisons, analyses across input configurations, and matched-indel controls were used to distinguish reproducible STR sensitivity from model- and sequence-design-specific effects. The resulting olive STR panel provides a genome-wide resource for population and trait analyses and demonstrates that STRs contribute complementary information about olive population structure. More broadly, this study evaluates how genomic foundation models represent natural multiallelic STRs and shows how embedding perturbation can complement phenotype association and genomic context in prioritizing candidate STRs for further investigation.

## Results

### Study Design

To comprehensively characterize short tandem repeat (STR) variation in olive (Olea europaea), we assembled a panel of 212 whole-genome sequenced accessions by integrating our in-house sequencing data with three publicly available datasets (Supplementary Data S1). The panel encompasses cultivated and wild olives sampled across the Mediterranean Basin and Iran, providing broad representation of the species’ genetic diversity. While previous population genomic and association studies in olive have primarily explored single-nucleotide polymorphisms (SNPs), this study focuses on genome-wide STR variation. We therefore genotyped STR loci in each accession using ConSTRain^30^, and after quality control and filtering, generated a genome-wide catalogue of polymorphic STRs (pSTRs) for downstream analyses.

The resulting pSTR catalogue provided the foundation for all subsequent analyses. We first characterized the genome-wide landscape of olive pSTRs, including their genomic distribution, motif composition, and genomic context. We next compared population genetic structure derived from genome-wide STRs and SNPs to evaluate the extent to which STR variation complements conventional SNP-based analyses in resolving genetic diversity. We then investigated whether plant genomic language models capture the effects of naturally occurring STR length variation on sequence representations by quantifying STR-induced embedding perturbation across three independently developed models. Finally, we integrated embedding perturbation with phenotype association analyses for fatty acid composition to prioritize candidate STRs supported by both sequence-based and phenotype-based evidence (Fig. 1).

**Figure 1:**
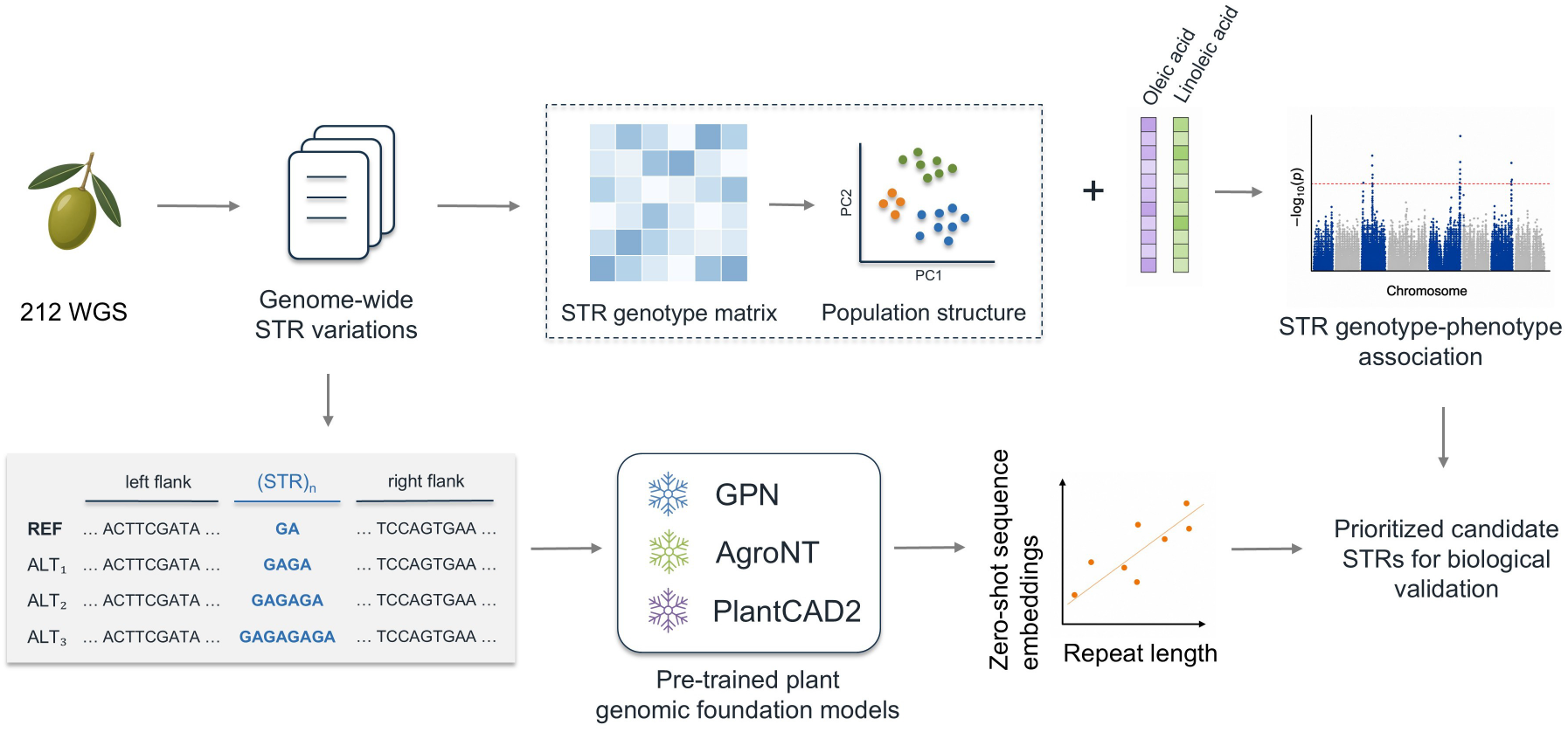
Framework for genome-wide functional interpretation of STR variation using plant genomic foundation models. Whole-genome sequencing data from 212 olive accessions were used to catalogue STR variation and construct a population genotype matrix. STR genotypes were analyzed independently for population structure and jointly with oleic- and linoleic-acid measurements for association mapping. Naturally observed STR alleles were also embedded with AgroNT, PlantCAD2, and GPN. Reference-anchored embedding perturbations were related to repeat-length differences and integrated with STR–phenotype evidence to prioritize candidate loci.

### Genome-wide landscape of STR variations in Olive

To define the STR variation available for downstream population and functional analyses, we first characterized the abundance, variability, and genomic distribution of STRs across the olive panel. After filtering, 117,119 polymorphic loci from 212 accessions were retained (Supplementary Data S3). The density of 65,095 chromosomal loci was 125.0 per megabase, approximately 1.9-fold higher than the density observed on unplaced scaffolds, where 52,024 loci corresponded to 65.4 per megabase (Fig. S1). Mononucleotide repeats dominated the catalogue, accounting for 59.0% of pSTRs, whereas the remaining motif classes each contributed substantially smaller proportions (Fig. 2A). Repeat-length variance differed markedly among motif classes: mononucleotide and dinucleotide loci showed the broadest distributions, whereas longer motifs were increasingly concentrated at lower variance (Fig. 2B). Therefore, Olive STR diversity is strongly shaped by motif length, with short-motif repeats comprising both the most abundant and the most variable components of the pSTR catalogue.

**Figure 2:**
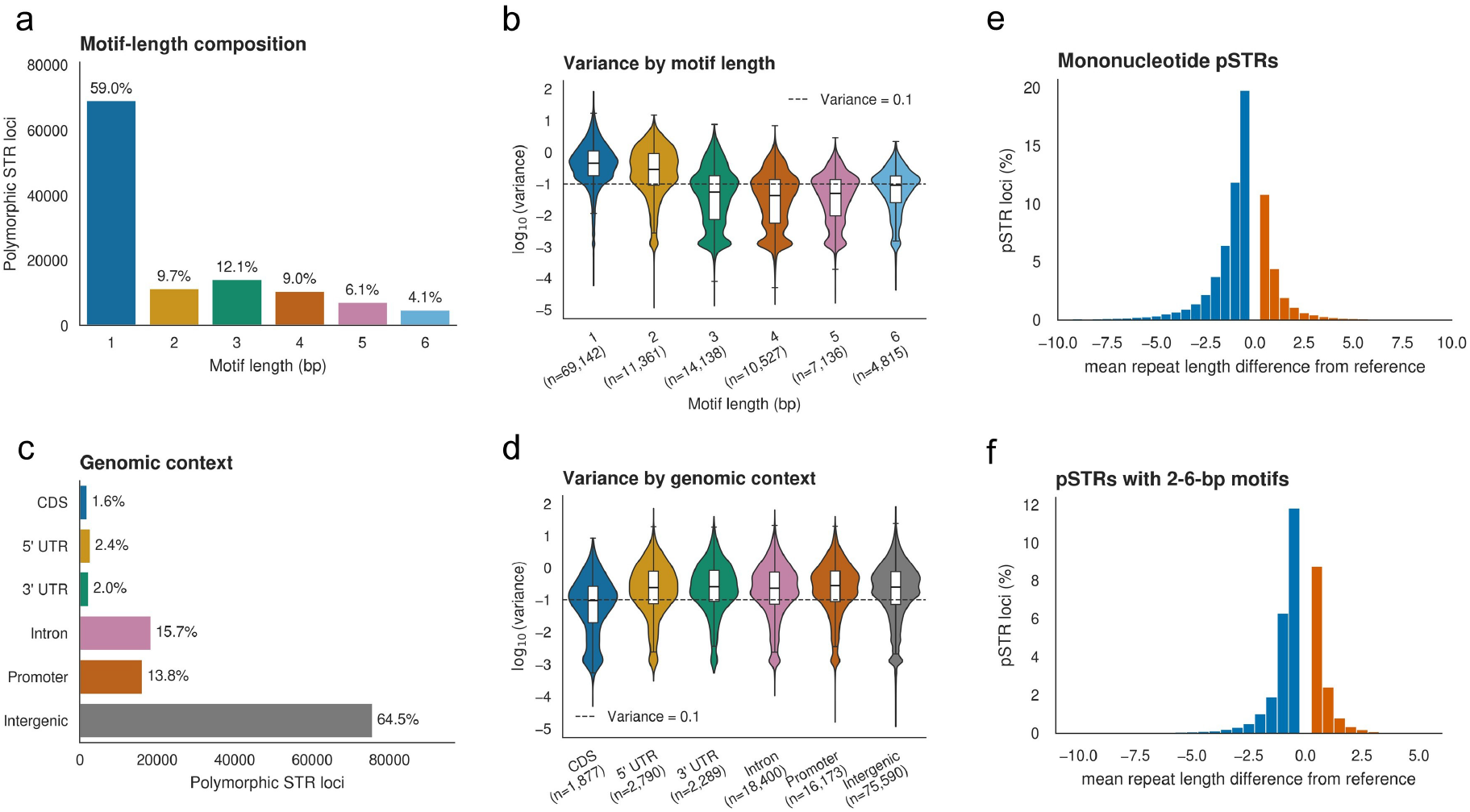
Genome-wide landscape of polymorphic STR variation in olive. **(A)** Counts and percentages of 117,119 polymorphic STRs by motif length. **(B)** Distribution of the base-10 logarithm of population repeat-length variance by motif length; the dashed line marks variance = 0.1. **(C)** Counts and percentages of polymorphic STRs by mutually exclusive genomic context; promoters are strand-aware regions 2 kb upstream of annotated transcription start sites. **(D)** Repeat-length variance distributions by genomic context. Violin interiors show boxplots. **(E,F)** Population-mean repeat-length difference from the reference allele for mononucleotide loci and loci with 2–6-bp motifs, respectively. Each locus contributes once; blue and orange bins denote negative and positive reference-relative differences, and the gray bin is centered on zero.

We next examined whether pSTR distribution and variability differed across genomic contexts. Most pSTRs were intergenic (64.5%), although more than one-third occurred within genes or their 2-kb promoter regions, particularly in introns and promoters (Fig. 2C). Variance distributions were broadly similar across genomic contexts, while coding STRs were shifted toward lower variation relative to noncoding classes (Fig. 2D), consistent with stronger constraint on length changes that may alter coding sequence. Motif composition showed different patterns in genomic context. Particularly, dinucleotide pSTRs were most strongly enriched in 5^′^ UTRs (1.96-fold; FDR < 0.05; Fig. S2). The presence of polymorphic repeats in promoters, UTRs, introns, and coding regions defines a subset of loci for which length variation could plausibly influence regulatory or transcribed sequence properties. The majority of repeat alleles differ in length from the reference genome by only a small number of repeat units, with a slight bias toward alleles that are shorter than the reference allele, and exhibit the broad distribution of variation at mononucleotide loci (Fig. 2E,F). This pattern may reflect greater biological variability of mononucleotide repeats, but could also be influenced by the reduced genotyping accuracy of short-read sequencing in homopolymeric regions. These distributions describe reference-relative states rather than observed mutation events, but together reveal a large and compositionally heterogeneous reservoir of repeat-length variation for further analyses.

### STRs resolve fine-scale population structure beyond SNPs

Previous studies in humans have shown that genome-wide STR variations can resolve high-resolution population structure^23^. Whether that also applies to olive populations remains unclear. We therefore compared population structure inferred from genome-wide SNPs and STRs to test whether STRs recapitulate major ancestry axes, while providing additional information for fine-scale population structure. To address this, we performed principal component analysis (PCA) using 299,285 polymorphic SNPs and 82,210 variance-filtered pSTRs across the same 212 accessions (Supplementary Data S4), and compared the population structure recovered by the two marker classes. Both marker classes separated Iranian, Western Mediterranean, and Eastern Mediterranean accessions and displaced wild *sylvestris* accessions from the main cultivated *europaea* distribution (Fig. 3A). The first STR principal component explained 9.38% of the variance, almost twice the 4.76% explained by SNP PC1; STR PC2 explained 3.63%, compared with 4.00% for SNP PC2 (Fig. 3B). Restricting to 45,666 pSTRs on assembled chromosomes produced nearly identical leading components (absolute score correlations, 0.99–1.00), indicating that the pattern was not driven by STRs on unplaced scaffolds.

**Figure 3:**
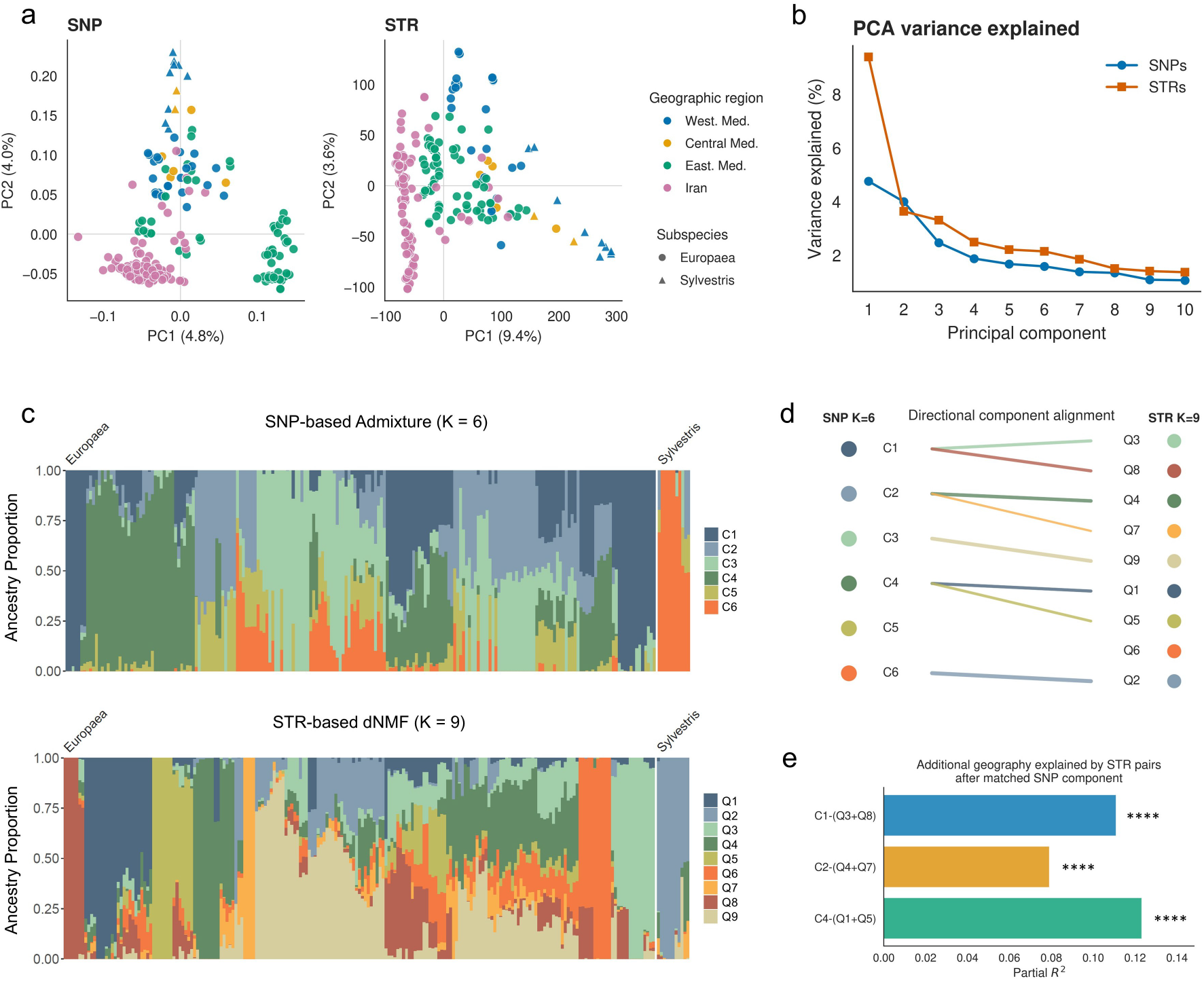
STRs recover shared ancestry and resolve additional fine-scale geographic structure. **(A)** SNP- and STR-based principal-component analyses of the same 212 accessions. Colors denote geographic region and shapes denote subspecies (See methods). **(B)** Variance explained by the first 10 SNP and STR principal components. **(C)** SNP ADMIXTURE at *K* = 6 and STR dNMF at *K* = 9, displayed in the same sample order. Accessions were ordered by hierarchical clustering of STR ancestry proportions using Jensen–Shannon distance within species. **(D)** Directional one-to-many alignment in which each qualifying STR component is assigned to its strongest SNP correlate; split SNP components connect to multiple STR components. **(E)** Additional geographic variance explained by each paired STR-component set after conditioning on its matched SNP component: Q3+Q8 conditioned on C1, Q4+Q7 on C2, and Q1+Q5 on C4. Bars show partial *R*^2^; four asterisks indicate permutation *P* = 0.0001 from 9,999 permutations.

We next examined whether the higher dimensionality of STR length variation translated into finer ancestry decomposition rather than simply reproducing the SNP-defined structure. To ensure a comparable genomic basis, both analyses were restricted to SNPs and STRs located on assembled chromosomes. We applied ADMIXTURE^31^ to SNPs and dNMF to STRs, and selected the optimal number of ancestry components according to the corresponding model-selection criterion. The results supported six SNP-based ancestry components (ADMIXTURE *K* = 6, minimum cross-validation error 0.3991) and nine STR-based ancestry components (dNMF *K* = 9; Fig. 3C and Fig. S3). For clarity, SNP-derived ancestry components are hereafter denoted C1–C6, whereas STR-derived components are denoted Q1–Q9. Both markers identified a distinct ancestry component to wild *sylvestris* accessions. To compare the more complex admixed structure within subsp. *europaea*, we matched every STR component to its strongest reproducible SNP correlate when that relationship passed the prespecified correlation and FDR thresholds. Three SNP ancestry components were each represented by a pair of STR components: Q3/Q8, Q4/Q7, and Q1/Q5. Summing each STR-component pair increased its correlation with the corresponding SNP component to 0.953, 0.887, and 0.937, respectively (Fig. 3D). By contrast, SNP component C5 and STR component Q6 had no qualifying match. This asymmetric correspondence indicates that STRs recover the major ancestry axes identified by SNPs while subdividing several of them into finer components. We then tested whether the three split STR-component pairs retained geographic information after conditioning on their matched SNP component. The Q3/Q8, Q4/Q7, and Q1/Q5 pairs explained an additional 11.1%, 7.9%, and 12.3% of multivariate geographic variation, respectively (all permutation *P* = 0.0001; Fig. 3E). Thus, STRs broadly reproduced the major ancestry structure inferred from SNPs while further resolving shared ancestry components in patterns associated with neighboring Mediterranean countries and Iranian provinces. These conditional gains support the interpretation that STR-based ancestry is complementary to, rather than simply a recapitulation of, SNP-based structure.

### Genome-wide quantification of STR-induced embedding perturbations

Recent genomic foundation models provide a powerful alternative approach for evaluating how sequence variation alters learned representations of DNA. Although these models were not trained specifically on olive, they have demonstrated the ability to estimate the potential functional consequences of sequence variants using zero-shot scores. We therefore examined whether naturally occurring STR length variation produced consistent embedding perturbations across three plant genomic language models: AgroNT, PlantCAD2, and GPN. For each locus, we measured the embedding distance between alternative alleles and the reference sequence and regressed this distance against the absolute difference in repeat length. The resulting slope, termed embedding sensitivity, quantified the magnitude of STR-induced embedding perturbation.

### Embedding perturbation across hidden layers and distance metrics

We examined whether naturally occurring STR length variation was encoded as a graded change in embedding distance from the reference allele. For each observed non-reference allele, we calculated the distance between its whole-sequence mean embedding and that of the corresponding reference allele and regressed this distance against the absolute difference in repeat count. The resulting slope was defined as embedding sensitivity. Statistical significance was assessed by permutation testing, followed by Benjamini–Hochberg correction, with loci at *q* < 0.05 classified as embedding-sensitive.

We first evaluated how STR-induced embedding perturbation varied across hidden layers and distance metrics to define the representations used in subsequent analyses. For AgroNT, layer 21 yielded the largest median whole-sequence slope among the four surveyed layers under both L1 and L2 distance, whereas the corresponding maximum for PlantCAD2 occurred at layer 36 (Fig. S4). Layer-wise patterns were broadly concordant between L1 and L2 distance, and the proportion of significant loci was similar across distance metrics within a given layer, although it varied across layers, particularly for PlantCAD2. We therefore selected AgroNT layer 21 and PlantCAD2 layer 36 for the primary analyses and used L2 distance because it directly quantifies the Euclidean distance between allele-specific embeddings. For GPN, we used the final hidden layer because a comparable layer-wise analysis was not available.

The common analysis set comprised 62,847 loci with at least three distinct non-reference repeat lengths (Supplementary Data S5). STR length variation produced a positive L2-distance slope at 98.1% of loci for AgroNT, 98.8% for PlantCAD2, and 96.5% for GPN (Fig. S6), indicating that greater repeat-length divergence was associated with progressively greater embedding perturbation across all three models.

### Cross-model reproducibility of embedding-sensitive loci

Using the primary representations and significance framework established above, we identified 12,296 embedding-sensitive loci in AgroNT, 12,662 in PlantCAD2, and 10,943 in GPN at *FDR* < 0.05 (Fig. 4A; Fig S5). An embedding perturbation detected by only one model could reflect architecture-specific behavior rather than a general property of the underlying sequence. We therefore assessed whether three independently trained plant genomic language models identified the same embedding-sensitive loci. Despite differences in model architecture and training objective and data, the three models converged on a large shared set of embedding-sensitive loci. A total of 7,417 loci passed FDR < 0.05 in all three models (Fig. 4B). Pairwise intersections comprised 9,911 loci for AgroNT–PlantCAD2, 8,828 for AgroNT–GPN, and 9,032 for PlantCAD2–GPN, corresponding to Jaccard indices of 0.66, 0.61, and 0.62 (Fig. 4C). Each overlap was at least fourfold greater than expected under independence (Fisher odds ratios, 58.3–72.2; all *P* < 10^−3^). Of the 10,943 GPN-significant loci, 10,443 (95.4%) were also significant in at least one other model, leaving only 500 loci specific to GPN. This convergence indicates that embedding sensitivity at these loci is not restricted to a single modeling framework.

**Figure 4:**
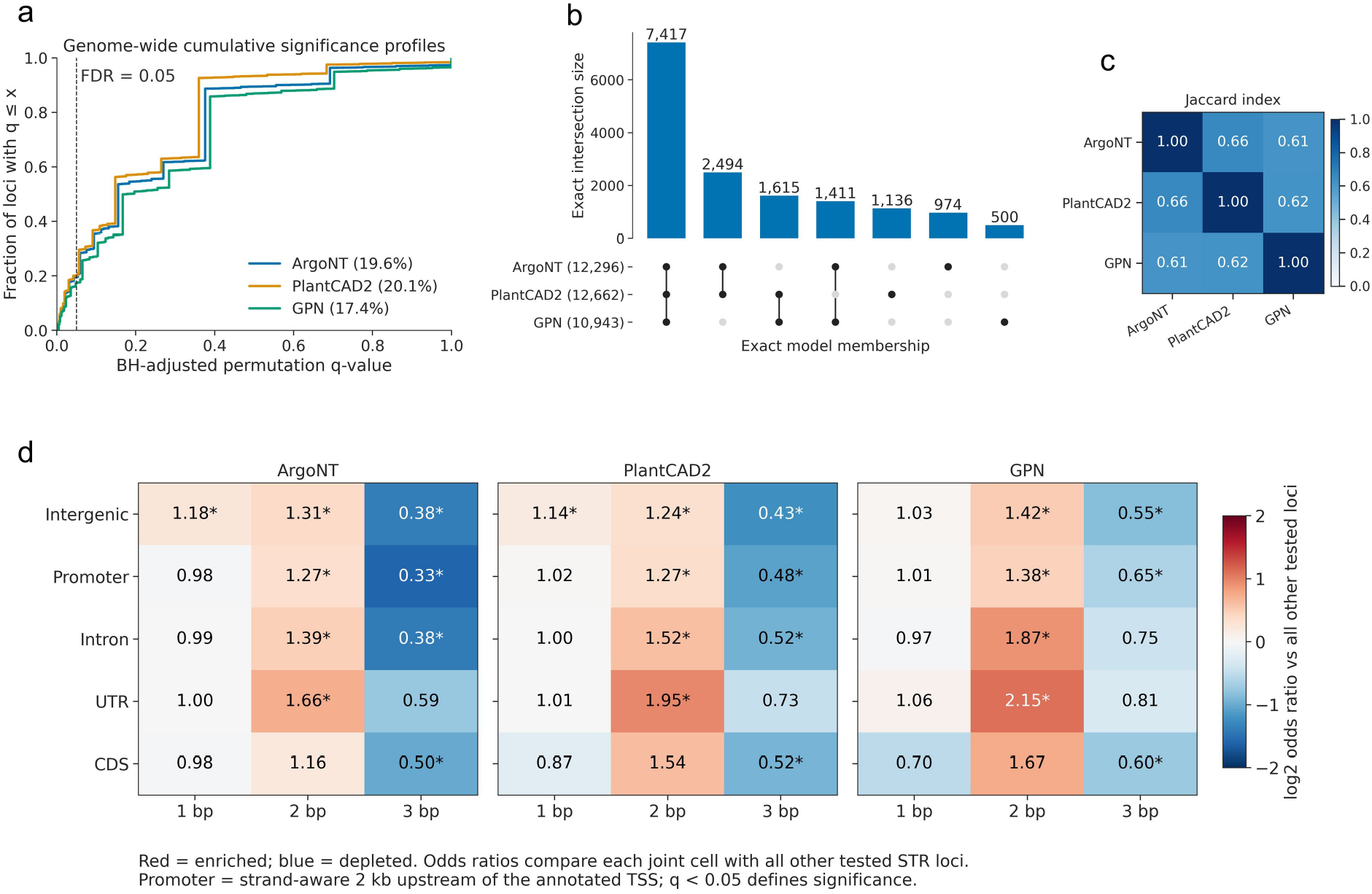
Genome-wide and cross-model reproducibility of STR-induced embedding perturbations. All panels compare the same 62,847 STR loci using reference-anchored whole-sequence mean L2 distances from AgroNT layer 21, PlantCAD2 layer 36, and the final GPN hidden layer. **(A)** Cumulative distributions of Benjamini–Hochberg-adjusted permutation *q* values. The dashed line marks *q* = 0.05; legend percentages are the fractions of tested loci below this threshold. **(B)** Exact intersections among model-specific significant sets. Bars give intersection sizes, and connected dots identify model membership. **(C)** Pairwise Jaccard indices for the significant-locus sets. **(D)** Enrichment or depletion of significant loci in joint motif-length–genomic-context categories. Cell values are odds ratios comparing each category with all other tested loci; colors show log_2_ odds ratios, and asterisks denote Benjamini–Hochberg FDR < 0.05. The UTR category combines 5^′^ and 3^′^ UTRs. Promoters are strand-aware 2-kb regions upstream of annotated transcription start sites.

Cross-model agreement was substantially stronger for significance calls than for the relative magnitude of embedding sensitivity. After robust standardization within each model, pairwise Pearson correlations of L2-slope magnitude ranged from 0.047 to 0.167, whereas Spearman correlations ranged from 0.064 to 0.217. Thus, the models reproducibly identified loci at which repeat-length variation altered sequence embeddings, but differed in how they ranked the magnitude of those changes. Consensus significance therefore represents the more transferable cross-model signal, whereas embedding-sensitivity magnitude retains substantial model-specific information.

### Sequence and genomic determinants of embedding sensitivity

To assess whether embedding sensitivity was associated with interpretable sequence and genomic features, we stratified loci by STR motif length and genomic context. Embedding-sensitive loci showed reproducible motif- and context-dependent patterns across all three models. Dinucleotide STRs were enriched among embedding-sensitive loci in every genomic context examined, with the strongest enrichment observed in UTRs and introns (odds ratios up to 2.15; Fig. 4D). By contrast, trinucleotide loci were consistently depleted, particularly in intergenic, promoter, and coding regions (odds ratios, 0.33–0.81). Enrichment of mononucleotide STRs was weaker and less consistent across models. Longer motif classes were excluded from these comparisons because of their limited representation within individual genomic contexts. The recurrence of the dinucleotide and trinucleotide patterns across models suggests that motif composition systematically influences whether STR length variation produces a statistically detectable embedding perturbation. The particularly strong enrichment of embedding-sensitive dinucleotide STRs in UTRs is consistent with the regulatory potential of repeat-length variation in transcribed non-coding regions.

The magnitude of embedding sensitivity also varied by motif length and genomic context. When normalized per base of repeat-length difference, AgroNT and PlantCAD2 generally showed larger responses for shorter motifs, whereas GPN showed relatively high sensitivity for mono- and tetranucleotide repeats. Across models, UTR, intronic, promoter, and intergenic STRs tended to exhibit greater per-base sensitivity than coding STRs, although their relative ordering differed among models (Fig. S7). Expressing sensitivity per repeat unit altered the motif-length pattern, as expected because longer motifs contribute more bases per added repeat, but broadly preserved the differences among genomic contexts (Fig. S8). Together, these results indicate that embedding sensitivity is structured by both repeat composition and genomic context, whereas model-specific differences in magnitude may reflect a combination of biological signal and technical factors, including differences in model architecture, training data, tokenization, and embedding representation.

### Robustness analyses of input design and matched-indel controls

Because statistical detection could potentially depend on the variation observed at each locus, we examined whether embedding-sensitive and nonsensitive loci differed in allelic diversity and population repeat-length variance. In all three models, embedding-sensitive loci occurred across a broad range of both measures, although they were somewhat more frequent among loci with greater allelic diversity and higher repeat-length variance (Fig. S6). Thus, the classification of embedding-sensitive loci was not restricted to highly variable STRs, indicating that observed allele diversity and repeat-length variation alone do not account for the detected embedding perturbations.

We next tested whether variation in total input length influenced the primary embedding estimates. We randomly sampled 1,000 eligible loci without replacement using a fixed random seed. This set was used for subsequent comparisons. Each allele was re-embedded using a fixed 500-bp left flank, with the right flank adjusted to produce a total input length of 1,000 bp. Agreement with the primary variable-length construction was high for AgroNT layer 21 (median relative L2 distance difference, 0.01; median Pearson correlation, 0.99) and PlantCAD2 layer 36 (0.08 and 0.99, respectively). GPN was not included because its primary inputs were already limited to 512 bp. These results indicate that the embedding patterns observed for AgroNT and PlantCAD2 were not primarily driven by STR-dependent variation in total input length.

Finally, to distinguish repeat-specific perturbation from generic effects of changing sequence length, we constructed length-matched nonrepetitive indel series from the local flanking sequence for the same subset of 1000 loci (Supplementary Data S7). STR embedding-sensitivity slopes exceeded the corresponding matched-indel mean at 92.0% of loci for AgroNT, 86.0% for Plant-CAD2, and 79.6% for GPN (paired sign-flip *P* = 10^−5^ for each model; Fig. S9). The mean per-base STR-minus-control differences were 0.423, 0.0236, and 0.370 for AgroNT, PlantCAD2, and GPN, respectively. Thus, changing sequence length contributes to the embedding response but is insufficient to explain its magnitude at most tested STRs, supporting a repeat-context-specific component.

### Prioritization of phenotype-associated STRs using embedding perturbation

Having identified embedding-sensitive STRs using three plant genomic foundation models, we next asked whether embedding perturbation could help prioritize loci associated with olive oil composition. Measurements of oleic and linoleic acid content were available for 140 accessions (Supplementary Data S2). The two traits were strongly inversely correlated (Pearson *r* = −0.919, *P* = 1.04 × 10^−5^^7^; Fig. S10), consistent with their closely related metabolic basis. We additionally derived the log-transformed oleic-to-linoleic acid ratio, log(oleic/linoleic), as a composite measure of fatty-acid composition. We tested 101,184 pSTRs with at least three observed allele lengths using trait-specific linear mixed models. Each model included measurement dataset and the first five STR principal components as fixed effects, with an SNP-derived kinship matrix included as a random effect. Six STRs reached Bonferroni-adjusted *P* < 0.05 in at least one trait-specific analysis (Fig. S11). Among these, the A/T mononucleotide STR at CACTIH010001827.1:963000 was the only locus associated with all three traits at the Bonferroni threshold (Fig. S12). For subsequent integration with embedding perturbation, we retained 68 unique STR loci that reached a within-trait Benjamini–Hochberg FDR of *q* < 0.05 in at least one analysis (Supplementary Data S6). These included 51 associations with oleic acid, 37 with linoleic acid, and 20 with the log-transformed ratio, with individual loci potentially associated with more than one trait. Nine loci reached *q* < 0.05 for all three phenotypes. Of the 68 unique phenotype-associated STRs, 45 were intergenic, 10 were located in promoters, 9 in introns, 3 in 5‘UTRs, and 1 in a coding region.

Of these 68 loci, 46 met the allele-count requirements for inclusion in all three embedding analyses. Among the eligible loci, seven were classified as embedding-sensitive in AgroNT, nine in PlantCAD2, and six in GPN (Fig. 5A). In total, 10 phenotype-associated STRs were supported by at least one model, including four that were embedding-sensitive in all three models.

**Figure 5:**
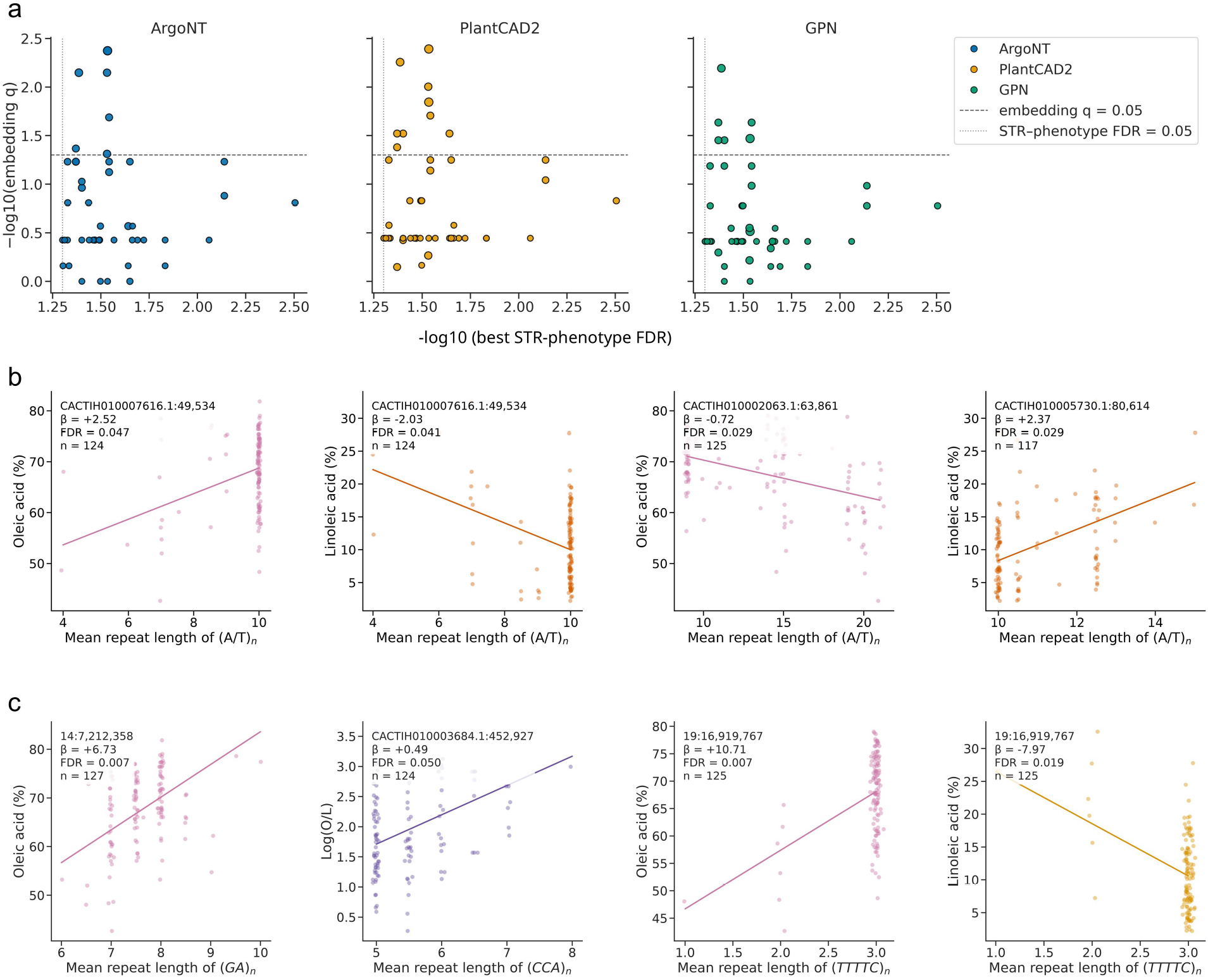
Integration of oil-trait associations with foundation-model embedding responses. **(A)** Best within-locus STR–phenotype FDR versus model-specific embedding FDR for phenotype-associated loci with matched embedding tests. Marker size is proportional to the number of observed nonreference alleles, and black outlines identify loci with embedding *q* < 0.05. Dashed horizontal and dotted vertical lines mark embedding and phenotype FDR = 0.05, respectively. **(B)** STR effects for three loci supported by all three embedding models and with concordant effect directions between the two phenotype measurement datasets. Points are raw sample measurements with horizontal jitter. Lines have the pooled covariate-adjusted linear mixed-model slope and are centered at the raw phenotype mean; annotations give the effect per additional repeat, phenotype FDR, and locus-wise sample size. CACTIH010007616.1:49,534 is shown for both oleic and linoleic acid. **(C)** Representative non-mononucleotide associations: the GA locus 14:7,212,358 with oleic acid, the CCA locus CACTIH010003684.1:452,927 with the log oleic/linoleic ratio, and the TTTTC locus 19:16,919,767 with oleic and linoleic ac3i0d. Points and lines are displayed as in panel B.

The four cross-model candidates were all A/T mononucleotide repeats, with embedding distance increasing as repeat length diverged from the reference allele (Fig. S13). One intronic STR at CACTIH010007616.1:49534 was associated with a 2.52-percentage-point increase in oleic acid content and a 2.03-percentage-point decrease in linoleic acid content per additional repeat unit (FDR = 0.047 and 0.041, respectively; Fig. 5B). An intergenic STR at CACTIH010002063.1:63861 was associated with lower oleic acid content (*β* = −0.72, FDR = 0.029), whereas a 5^′^UTR STR at CACTIH010005730.1:80614 was associated with higher linoleic acid content (*β* = 2.37, FDR = 0.029). The fourth candidate, an intergenic STR at CACTIH010005604.1:173698, was associated with lower linoleic acid content (*β* = −2.22, FDR = 0.043).

Although 50 of the 68 phenotype-associated STRs were mononucleotide repeats, non-mononucleotide loci provided complementary candidates with comparatively large estimated effects (Fig. 5C). Increasing GA repeat length at 14:7212358 was associated with higher oleic acid content (*β* = 6.73, FDR = 0.0073). The pentanucleotide STR at 19:16919767 showed opposing associations with oleic and linoleic acid content (*β* = 10.71, FDR = 0.0073, and *β* = −7.97, FDR = 0.019, respectively). A CCA repeat in the 5^′^UTR at CACTIH010003684.1:452927 was associated with a higher log-transformed oleic-to-linoleic acid ratio (*β* = 0.486, FDR = 0.050). These loci extend the candidate set across multiple repeat classes, although several effect estimates were supported by small minor repeat-length groups and should therefore be interpreted cautiously.

Together, these results indicate that embedding perturbation provides a complementary sequence-based criterion for prioritizing phenotype-associated STRs. Cross-model consensus highlighted a small subset of reproducible candidates, while trait-associated loci from additional repeat classes broadened the candidate set. We therefore interpret the joint phenotype-association and embedding evidence as a framework for candidate prioritization and experimental follow-up, rather than as proof of causal effects on fatty-acid composition.

## Discussion

This study integrates three aspects of short tandem repeat (STR) variation that are commonly examined separately: genome-wide diversity, population structure, and sequence-based interpretation. We generated a catalogue of 117,119 polymorphic STRs in olive and showed that genome-wide STRs recover the major ancestry patterns inferred from SNPs while resolving additional geographically structured variation. We further evaluated how three plant genomic foundation models represent naturally occurring multiallelic STR series and integrated reproducible embedding perturbation with oil-composition associations to prioritize candidate loci. Together, these analyses demonstrate the potential of leveraging zero-shot representations from state-of-the-art plant genomic language models to support the interpretation and prioritization of naturally occurring STR variation.

The olive STR landscape was strongly structured by motif composition and genomic context. Mononucleotide repeats were the most abundant polymorphic class, whereas mono- and dinucleotide STRs generally displayed broader repeat-length variation than longer motifs, consistent with established effects of motif architecture on STR mutation dynamics^13,32^. Coding STRs were less variable than noncoding loci, compatible with stronger constraint on length-changing alleles in protein-coding regions^33,34^. These patterns nevertheless reflect observable variation under the OLEA9 reference and the applied filtering scheme. Short repeats remain particularly susceptible to sequencing and genotyping error^35^, and differences between assembled chromosomes and unplaced scaffolds may reflect assembly quality and mappability as well as biology. Likewise, the excess of population-mean contractions relative to OLEA9 cannot be interpreted as a directional mutational bias without ancestral polarization.

STRs and SNPs captured the same broad ancestry pattern but differed in their resolution of recent population differentiation. Both marker classes separated Iranian, Eastern Mediterranean, Western Mediterranean, and wild ancestry, whereas STRs subdivided several SNP-defined components. These subdivisions retained geographic information after conditioning on the corresponding SNP ancestry component, indicating that they were not simply redundant decompositions of the same signal. The faster mutation rate and multiallelic state space of STRs may preserve aspects of recent differentiation that are represented less distinctly by linked SNPs^21,23,36^. This extends previous olive studies based on small microsatellite panels to a genome-wide setting^19^. Beyond the present analyses, the genome-wide STR catalogue provides a reference for population genetic studies, germplasm characterization, parentage and cultivar identification, association mapping, and the prioritization of repeat variants for breeding-related traits. Its integration with SNP data may be particularly useful for resolving closely related accessions and identifying fine-scale structure that could otherwise remain undetected.

Natural STR allelic series provide a distinct test for genomic foundation models because repeat-length changes simultaneously alter motif dosage, sequence length, and spacing between flanking elements. Across AgroNT, PlantCAD2, and GPN, embedding distance generally increased with absolute repeat-length difference from the reference allele, demonstrating that all three models encoded many STR series as graded changes rather than simply distinguishing reference from non-reference sequence. Approximately one fifth of tested loci were classified as embedding-sensitive, and 7,417 were significant in all three models. Pairwise overlaps greatly exceeded independence expectations, indicating that sensitivity at many loci was not confined to one architecture or training corpus. By contrast, correlations between standardized slope magnitudes were weak. The models therefore agreed more strongly on whether an STR altered their learned representation than on the relative magnitude of that change. Because the models differ in training data, objectives, context windows, sequence representation, and hidden-state geometry^26–28^, consensus significance is a more defensible cross-model prioritization criterion than direct comparison of raw embedding distances.

Embedding sensitivity was also associated with repeat composition and genomic context. Dinucleotide STRs were repeatedly enriched among embedding-sensitive loci, whereas trinucleotide STRs were depleted across several genomic contexts. Such patterns are compatible with known relationships between STR composition, genomic location, and regulatory sequence properties^15,37,38^. The recurrence of these patterns across independently trained models suggests that their internal representations may have learned sequence features linked to genomic regulation and local sequence architecture, rather than responding only to changes in repeat length. This interpretation is further supported by the robustness analyses: variation in total input length was not the primary source of the AgroNT and PlantCAD2 signals, and embedding sensitivity at STR loci exceeded those of length-matched nonrepetitive indels at most tested loci in all three models. Together, these findings raise the possibility that plant genomic foundation models have implicitly learned regulatory and structural sequence features that shape how STR length variation is represented.

The oil-composition analysis illustrates how cross-model embedding evidence can complement phenotype-association results. Oleic and linoleic acid were strongly inversely correlated, consistent with their connected biosynthetic pathways and previous genetic studies of olive oil composition^7–9^. Four phenotype-associated STRs were embedding-sensitive in all three models, providing convergent population-level and sequence-representation evidence. Additional non-mononucleotide loci broadened the candidate set across repeat classes. However, none of the four cross-model candidates passed the trait-wise Bonferroni threshold. Association and embedding analyses therefore provide distinct but complementary information: association tests identify loci whose repeat lengths covary with phenotype in the sampled population, whereas embedding analyses identify allelic changes that are represented reproducibly across independently trained models. Their intersection prioritizes loci supported by two independent lines of evidence.

Several limitations define the next steps. Short-read STR genotyping remains challenging for long, interrupted, and structurally complex repeats, and reliance on a single linear reference may introduce mapping and reference-allele biases^35,39^. Long-read sequencing and an olive pangenome should improve allele resolution and identify sequence configurations absent from OLEA9. The phenotype analysis was limited to 140 accessions, reducing power for rare alleles and limiting the separation of genetic, environmental, and dataset effects. Independent population and environmental replication will therefore be essential. In addition, the genomic foundation models were not trained specifically on olive sequences, so their representations may not capture species-specific regulatory grammar or repeat architecture. Future models trained or adapted using olive and related Oleaceae genomes may provide more accurate and biologically relevant representations. Finally, the zero-shot embeddings were not accompanied by expression, chromatin, or biochemical measurements. Reporter assays, allele-specific expression, chromatin profiling, and targeted repeat editing will be needed to determine which prioritized STRs alter molecular phenotypes and oil composition.

In summary, this study provides a genome-wide STR resource for olive and demonstrates that STRs contribute to population-structure information beyond SNPs alone. It also establishes a general framework for evaluating how independently trained genomic foundation models represent natural multiallelic STR variation while separating reproducible repeat-context sensitivity from generic sequence-length effects. Integrating this evidence with population genetics and phenotype association narrows the candidate space for experimental investigation without substituting computational predictions for validation. Applied to larger cohorts, pangenomes, and functional datasets, this framework could help translate repeat variation into testable biological and breeding-relevant hypotheses in olive and other crops^11,29^.

## Methods

### Plant material and genomic datasets

The study panel comprised 212 olive (Olea europaea) accessions assembled from three publicly available whole-genome sequencing datasets and one previously unpublished dataset. The public component comprised 107 genomes, including 51 accessions from BioProject PRJNA556567, 43 from PRJNA782823, and 12 from PRJEB35540^6,10,40^. EU50 (PRJNA556567) was excluded because its BAM file failed integrity validation, leaving 105 public accessions. The remaining 107 accessions from the Iranian cohort consisted of previously unpublished whole-genome sequencing data generated by the co-authors of the present study and included here with permission (accession numbers PRJNA1542630 and PRJNA1542632, which are currently not publicly accessible). Detailed descriptions of sample collection, library preparation, sequencing platforms, and primary read preprocessing will be reported separately. These accessions were maintained by the Iran Olive Germplasm Bank of Minoodasht and sampled across 12 provinces. In total, there are 212 accessions analysed in this study: 201 subsp. *europaea* and 11 subsp. *sylvestris*. All genome-coordinate analyses used the soft-masked *O. europaea* OLEA9 assembly and Ensembl Plants release 60 annotation^3^.

### Oil-quality phenotyping and collection

Measurements of oleic and linoleic acids were collected from 29, 9, and 9 accessions of BioProjects PRJNA556567, PRJEB35540, and PRJNA782823. For the 93 accessions in the Iranian cohort, olive oil was obtained by cold pressing and centrifugation at 4 ^◦^C. Fatty-acid methyl esters were quantified with a Younglin 6100A gas chromatograph equipped with an HP-88 column, and fatty acids were identified following International Olive Council procedures^41^. The analyses used oleic-acid percentage, linoleic-acid percentage, and the natural logarithm of their ratio.

### STR genotyping and quality control

STR genotyping was implemented as a DSL2 Nextflow workflow^42^. The target catalogue containing 480,768 STRs with 1–6-bp motifs was generated using mreps^43^ on the OLEA9 assembly. Control-FREEC v11.6b^44^ was run separately for each coordinate-sorted BAM under a diploid baseline, using 5-kb read-depth windows. Segmented calls were converted to ConSTRain BED input after retaining integer absolute copy numbers from 0 to 20 that differed from the baseline and spanned at least 10 kb. Each sample was then processed once with the CNV-aware mode of ConSTRain v1.1.0^30^. Only PASS genotypes were retained. At each locus, the arithmetic mean of the reported allele-copy repeat lengths represented an accession’s genotype. Loci were required to have calls in at least 80% of the 212 accessions and more than one distinct non-missing length. This produced 117,119 polymorphic STRs (pSTRs) for the genome-wide landscape analyses. 82,210 loci with variance ≥ 0.1 were retained for primary STR PCA, whereas later analyses applied their stated allele-number or strict-variance requirements.

### Genomic annotation and STR landscape analyses

The complete target catalogue and the 117,119 pSTRs were intersected with Ensembl Plants release 60 gene models using BEDTools^45^. Mutually exclusive annotations were assigned in the hierarchy CDS, 5^′^ UTR, 3^′^ UTR, other exon, intron, promoter, and intergenic. Promoters were defined as the strand-aware 2-kb interval immediately upstream of an annotated gene boundary and were assigned only after all genic categories. We summarized pSTR counts, repeat-length variance, motif class, genomic context, and chromosome or unplaced-scaffold density. Reference repeat number was calculated as target length divided by motif length; for each pSTR, the population mean repeat number minus the reference value was summarized in 0.5-repeat bins. Motif-by-context enrichment compared the pSTR fraction with that in the complete 480,768-locus catalogue using two-sided Fisher exact tests, with Benjamini–Hochberg correction across the 36 motif–context tests^46^.

### SNP- and STR-based population structure

SNP genotypes were generated by co-authors using a GATK-based diploid variant-calling pipeline. Full details of primary variant discovery will be reported separately. For the present analyses, the genotype dataset was first restricted to the 212 study accessions. The filtered SNP set was then processed using PLINK v1.9.0-b.7.7 with filters for variant missingness, sample missingness, and minor-allele frequency (--geno 0.05 --mind 0.05 --maf 0.05), followed by linkage-disequilibrium pruning (--indep-pairwise 50 10 0.2). This yielded 299,285 SNPs across the 23 assembled olive chromosomes for principal-component analysis and kinship estimation. Population structure was compared in the 212 accessions shared by the SNP and STR matrices. For STR PCA, missing values at the 82,210 variance-filtered loci were imputed with the locus mean, after which loci were centered and scaled to unit variance. A sensitivity PCA used the 45,666 variance-filtered STRs assigned to chromosomes 1–23. For visualization, country-of-origin annotations were aggregated into three broad geographic regions: Western Mediterranean (Spain, Morocco and Algeria), Central Mediterranean–Adriatic (Italy, Tunisia, Croatia and Albania), Eastern Mediterranean–Levant (Greece, Turkey, Syria, Lebanon and Israel). The same chromosomal STR set was then used for STR-based ancestry proportion estimation. SNP-based ancestry proportions were estimated with ADMIXTURE for *K* = 3–15, and the rank with the minimum cross-validation error was selected^31^. STR-based ancestry proportions were estimated using dNMF for *k*3–15, and the rank with maximum stable components was selected^23^. The ancestry coefficients from the selected positive channel were used downstream.

### Directional one-to-many ancestry-component alignment

To avoid forcing correspondence between marker systems to be one-to-one, each of the nine STR components was assigned to the SNP component with which it had the largest positive Pearson correlation. An assignment was retained only when *r* ≥ 0.30 and Benjamini–Hochberg FDR ≤ 0.05 both in the complete cohort and after restricting to the 201 subsp. *europaea* accessions. For SNP components receiving multiple STR assignments, the assigned STR proportions were summed and their correlation with the SNP component was compared with that of the strongest individual STR component. The improvement and its 95% interval were evaluated with 9,999 sample-level bootstrap replicates.

We next tested whether the three observed component splits: C1→Q3/Q8, C2→Q4/Q7, and C4→Q1/Q5, contained geographic information beyond their matched SNP component. The analysis retained 185 subsp. *europaea* accessions from countries or Iranian provinces represented by at least five accessions. Fine-geographic categories were encoded by orthonormal contrasts. For each split, a reduced multivariate model containing the SNP component was compared with a full model that additionally contained both assigned STR components. Partial *R*^2^ quantified the additional geographic variance, and significance was assessed with 9,999 Freedman–Lane residual permutations^47^.

### STR allele reconstruction and sequence-model embedding

The embedding catalogue comprised 75,802 loci with at least three distinct repeat counts among the variance-filtered pSTRs. The 394,921 embedded allele sequences represented unique locus–repeat-count combinations rather than individual-accession genotypes. Of these, 394,845 combinations were observed among the called genotypes, and 76 additional OLEA9 reference-allele combinations absent from the panel were added to provide a reference anchor at every locus. Thus, the OLEA9 reference allele was embedded alongside all observed non-reference alleles. Each allele was reconstructed by replacing the OLEA9 reference repeat tract with the corresponding number of motif copies while retaining the reference flanking sequence. Sequence characters were converted to uppercase, and all characters other than A, C, G, or T were normalized to N. AgroNT retained N as its ambiguity token, whereas the base-level PlantCAD2 and GPN tokenizers mapped N to [UNK]. Both models received 500 bp of left reference flank, the allele-specific STR, and 500 bp of right reference flank, such that total input length varied with allele length. AgroNT embeddings were extracted from layers 11, 21, 31, and 40 (1,500 dimensions) and averaged across six tokenizer frames and forward and reverse-complement orientations. PlantCAD2 embeddings were extracted from layers 12, 24, 36, and 48 (1,536 dimensions) after combining the reverse-complement parameter-sharing channels.

The 512-bp input of GPN contained the final 256 bp of left flank, the complete allele-specific STR, and sufficient right-flanking sequence to reach a total length of 512 bp. Forward- and reverse-complement token embeddings were aligned by genomic coordinate and averaged, and the 512-dimensional final hidden state from convolutional block 25 was retained. No tokenizer-level truncation was applied. AgroNT inputs were screened to fit within 1,026 token positions, PlantCAD2 received each complete reconstructed sequence, and GPN reconstruction raised an error if the full STR tract could not be accommodated within the 512-bp design. All retained alleles satisfied these criteria.

For all models, we calculated both the mean embedding over the complete biological input and the mean embedding over the allele-specific STR tract. Padding and special-token positions were excluded from whole-input pooling. For AgroNT, the [CLS] position and batch padding were omitted using the sequence length and attention mask; PlantCAD2 and GPN were run without inserted special tokens or padding.

For AgroNT, tokenizer outputs were mapped back to nucleotide intervals, and STR pooling included every token whose interval overlapped the allele-specific repeat tract, including tokens spanning a tract boundary. Coordinates were transformed for reverse-complement orientations and shifted for each tokenizer frame. PlantCAD2 and GPN used one token per nucleotide, allowing direct mapping of the STR interval to token positions. Zero-repeat alleles were represented at the deletion junction: AgroNT used the token spanning the junction, or both adjacent tokens when the junction coincided with a token boundary, whereas PlantCAD2 and GPN used the two nucleotide positions flanking the junction.

AgroNT^26^, PlantCAD2-Large^28^, and GPN^24^ were obtained from Hugging Face using the model repositories InstaDeepAI/agro-nucleotide-transformer-1b, kuleshov-group/PlantCAD2-Large-l48 and songlab/gpn-brassicales, respectively. All models were run in evaluation mode without gradient computation or stochastic sampling. AgroNT and PlantCAD2 inference used float16 and bfloat16 precision, respectively, whereas GPN used float32. Hidden states were converted to float32 before strand or frame combination and pooling, and all resulting embeddings were stored in float32. Because inference included no intentional stochastic operation, no random seed was applied. Embedding inference was performed on NVIDIA A100 GPUs.

### Reference-anchored embedding perturbation analysis

The catalogue-level inclusion criterion required at least three distinct observed repeat counts per locus and could therefore count the reference allele among those three. The reference-anchored regression instead treated the reference only as the embedding-distance anchor and required at least three distinct observed non-reference repeat counts as regression observations. Of the 75,802 embedded loci, 12,955 had exactly two observed non-reference repeat counts because their reference repeat count was also observed; these loci were excluded because a linear model with an intercept fitted to only two observations is saturated and does not support informative permutation inference. The canonical analysis therefore retained 62,847 loci. For every observed non-reference allele, the predictor was the absolute repeat-count difference from the reference and the response was its L1, L2, or cosine distance from the reference embedding. An intercept was included. Allele labels were enumerated exactly when the number of permutations was at most 10,000; otherwise 10,000 deterministic Monte Carlo permutations were used to obtain two-sided locus-level *P* values. Benjamini–Hochberg correction was applied separately within each pooling, layer, and distance family.

Layer selection was performed before cross-model comparison by examining the median genome-wide slope and significant fraction across the surveyed layers and metrics (Fig. S4). AgroNT layer 21 and PlantCAD2 layer 36 gave the strongest median L1/L2 responses and were selected; GPN contributed its single final hidden layer. Whole-sequence mean pooling and L2 distance were designated the primary common setting because L2 measures the Euclidean magnitude of the embedding displacement. L1 and cosine distance, alternative layers, and STR-tract pooling were retained as sensitivity analyses.

### Cross-model reproducibility and genomic determinants

Cross-model comparison used the same 62,847 loci and the stored primary permutation FDR values. Model-specific significant sets at FDR < 0.05 were compared by exact intersections and Jaccard indices. Pairwise overlap was tested against the common locus universe with one-sided Fisher exact and hypergeometric tests. Because raw distances are architecture- and layer-dependent, magnitude comparisons used the absolute L2 slope per base, followed by log_10_ transformation and within-model robust standardization as (*x* − median(*x*))/(1.4826 MAD(*x*)). Pearson and Spearman correlations compared these normalized magnitudes. Motif-length and genomic-context associations with significance were tested with Fisher exact tests and Benjamini–Hochberg correction; significant-locus slopes were also summarized per base and per repeat.

### Input-length robustness and matched-indel controls

Input-length robustness was evaluated in a randomly selected subset of 1,000 loci containing 5,697 alleles. AgroNT and PlantCAD2 alleles were re-embedded using exactly 500 bp of left flank and sufficient right flank to make every input 1,000 bp long. Each fixed-total embedding was compared directly with the corresponding primary variable-total embedding using cosine similarity, relative L2 difference, mean absolute difference, and Pearson correlation; no regression or significance test was applied. The comparison used AgroNT layer 21 and PlantCAD2 layer 36. GPN was omitted because its primary input was already fixed at 512 bp.

To separate repeat-specific effects from generic length changes, we constructed three deterministic matched-indel replicates for each of the same 1,000 loci. A/C/G/T bases from the local non-STR flanks were pooled and shuffled with locus- and replicate-specific seeds; candidate tapes containing at least four tandem copies of any 1–6-bp motif were rejected. Control alleles were nested prefixes whose lengths exactly matched the corresponding STR alleles. Each model retained its primary context construction, and control distances were anchored to the length-matched control reference. L2 slopes per repeat and per base were averaged across the three controls and compared with the genuine STR slope. Uncertainty was estimated with 10,000 locus bootstraps and significance with 100,000 paired sign flips.

### STR association analyses of oil-quality traits

Association analyses included all 140 phenotyped accessions. The STR genotype was the mean repeat length across the allele copies reported for an accession, 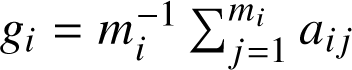, where *a_ij_* is the repeat count of allele copy *j* ; thus, for an ordinary diploid genotype, *g_i_* = (*a_i_*_1_ + *a_i_*_2_)/2. Of the 117,119 pSTRs, 101,184 had at least three distinct observed mean repeat lengths across the combined phenotype cohort and were tested; no additional call-rate threshold was imposed. Missing STR genotypes were excluded independently at each locus without imputation, producing locus-wise sample sizes of 104–140. Oleic-acid percentage, linoleic-acid percentage, and log(oleic/linoleic) were analyzed. No phenotype observations were excluded as outliers and no rank-based normalization was applied. Oleic and linoleic acid were analyzed on their original percentage scales, whereas their ratio was natural-log transformed. Model assessment used null-model convergence and variance-component summaries, quantile–quantile plots, and genomic-inflation factors; no formal residual-normality test or locus-specific residual diagnostic was applied.

The primary model was a linear mixed model with STR mean repeat length, phenotype measurement dataset, and STR PCs 1–5 as fixed effects and a genome-wide SNP-derived kinship matrix as a random effect^48^. STR PCs were calculated once across the full panel of 212 accessions from all 82,210 pSTRs that passed the population-variance threshold of 0.1. At each locus, missing genotypes were replaced by the locus mean and the locus was centered and scaled by its sample standard deviation before eigendecomposition; PC scores were then subset to the phenotyped accessions, and PCs 1–5 were standardized within each analysis cohort before inclusion as covariates.

The SNP kinship matrix was derived from 299,285 chromosome SNPs. The input panel contained biallelic A/C/G/T SNPs passing genotype missingness ≤ 0.05 and minor-allele frequency ≥ 0.05, followed by linkage-disequilibrium pruning. PLINK v1.9.0-b.7.7^49^ was used to compute all 212 principal components and eigenvalues in the accessions shared by the SNP and STR datasets. The genomic relationship matrix was reconstructed as *K* = *U*Λ*U*^T^ from the complete eigenvector matrix *U* and eigenvalue matrix Λ, symmetrized, scaled to a mean diagonal of one, and subset to the phenotyped accessions.

Associations were fitted with a custom Python 3.10.20 implementation using NumPy 2.2.6, pandas 2.3.3, and SciPy 1.15.2. For each phenotype and analysis cohort, the genetic-to-residual variance ratio was estimated once under the covariate-only null model by restricted maximum likelihood and then reused for all 101,184 loci in that trait–cohort scan, following the EMMAX/P3D approximation^48^. At each locus, the STR coefficient and standard error were nevertheless refitted by generalized least squares using the locus-specific complete cases and corresponding kinship submatrix. For oleic and linoleic acid, the effect estimate represents the change in percentage points per one-repeat increase in mean repeat length; for log(oleic/linoleic), it represents the change in log ratio per additional repeat, with 100[exp( *β*) − 1] giving the corresponding percentage change in the ratio. Formal discoveries were defined separately for each phenotype using the Bonferroni threshold 0.05/101,184 = 4.9415 × 10^−7^. Exploratory associations were defined as Benjamini–Hochberg *q* < 0.05, with adjustment performed independently within each phenotype across the 101,184 tests.

### Integration of association and embedding evidence

The exploratory association follow-up contained 108 phenotype–locus associations at FDR < 0.05, representing 68 unique loci. Of these, 46 satisfied the alternate-allele requirements of the matched three-model embedding analysis. For each locus we recorded model-specific embedding FDR and the number of models supporting an embedding response; significance in all three models defined cross-model consensus support. Candidate plots retained the pooled covariate-adjusted mixed-model effect, dataset-specific effect direction, genomic annotation, and the reference-anchored allelic embedding series. These analyses were used for prioritization rather than as evidence that an associated STR was causal.

## Acknowledgments

The computational work in this study was performed on the Earth cluster operated by the HPC team at the School of Life Sciences and Facility Management, Zurich University of Applied Sciences. We would like to thank Dr. Max Verbiest for his support in the initial construction of the olive STR panel.

## Author contributions

F.X. and M.H.P. conceived the study. M.H.P. established the initial data-preprocessing workflow. F.X. developed the computational methodology, performed the analyses, prepared the figures, and wrote the original manuscript. M.H.P., M.H.M., A.A., Y.Z. and J.Z. contributed to olive accessions, phenotype measurements and interpretation of the results. M.A. and V.S. supervised the study and acquired funding. All authors reviewed, edited, and approved the final manuscript.

## Ethics, Consent to Participate, and Consent to Publish declarations

Not applicable.

## Data and code availability

The publicly available whole-genome sequencing data analyzed in this study are available under BioProject accessions PRJNA556567, PRJEB35540, and PR-JNA782823. The unpublished sequencing dataset generated by co-authors of the present study is not currently publicly available because it forms part of an ongoing study. Requests for access should be directed to the corresponding author and will be considered in consultation with the data generators. The OLEA9 reference genome assembly used in this study was obtained from Ensembl Plants release 60. The soft-masked top-level sequence is available as Olea_europaea.OLEA9.dna_sm.toplevel.fa.gz. The Nextflow workflow used for STR genotyping is available at https://github.com/Garfieldfei/olive_str_genotyping. All remaining analysis and figure-generation code is available at https://github.com/acg-team/Olive_STRs.

## Supplementary materials

Supplementary Fig. S1 to S13

Supplementary Data S1 to S7

