## Supplementary Data S1-S7 for "Interpreting Short Tandem Repeat Variations with Genomic Foundation Models in Olive": README.md

# Supplementary data

## Supplementary Data S1 — Sample manifest

`Supplementary_Data_S1.csv` contains sequencing, mapping, coverage, geographic, taxonomic, source, and STR callability information for the 212 retained olive accessions. EU50 is excluded.

## Supplementary Data S2 — Oil-composition phenotypes

`Supplementary_Data_S2.csv` contains oleic- and linoleic-acid measurements, their ratio, and the natural-log-transformed ratio for the 140 phenotyped accessions.

## Supplementary Data S3 — Polymorphic STR catalogue

`Supplementary_Data_S3.tsv.gz` is a gzip-compressed catalogue of 117,119 pSTRs, including OLEA9 coordinates, repeat motifs, population variation and callability statistics, and functional annotations.

## Supplementary Data S4 — Population structure

`Supplementary_Data_S4.tsv` contains SNP and STR PC1/PC2 scores and the selected SNP ADMIXTURE K=6 and STR dNMF K=9 ancestry proportions for all 212 retained accessions.

## Supplementary Data S5 — Primary embedding results

`Supplementary_Data_S5.tsv.gz` reports primary AgroNT, PlantCAD2, and GPN embedding slopes, permutation P values, FDR values, normalized magnitudes, and cross-model significance for 62,847 matched STR loci.

## Supplementary Data S6 — Exploratory STR associations

`Supplementary_Data_S6.csv` contains 108 phenotype associations representing 68 unique STRs that passed within-trait FDR < 0.05, together with effect estimates, locus annotations, cohort-level results, and embedding support.

## Supplementary Data S7 — Embedding robustness

`Supplementary_Data_S7.csv` summarizes the primary fixed-input-length and matched-nonrepetitive-indel controls for AgroNT, PlantCAD2, and GPN.
