## Supplementary Fig S1-S13 for "Interpreting Short Tandem Repeat Variations with Genomic Foundation Models in Olive"

### This PDF file includes:

Supplementary Figures S1 to S13

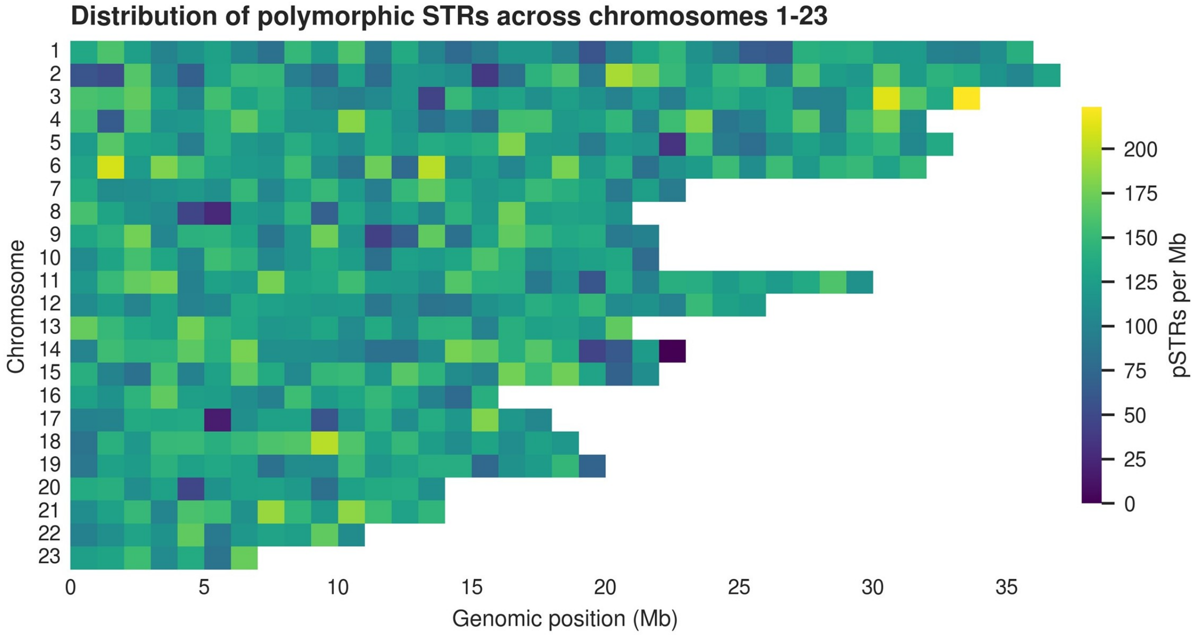


**Figure S1**: **Chromosomal distribution of polymorphic STRs.** Heatmap of pSTR density in 1-Mb windows across the 23 assembled olive chromosomes. Color represents the number of pSTRs per megabase; blank cells fall beyond the assembled length of the corresponding chromosome. Unplaced scaffolds are not displayed.

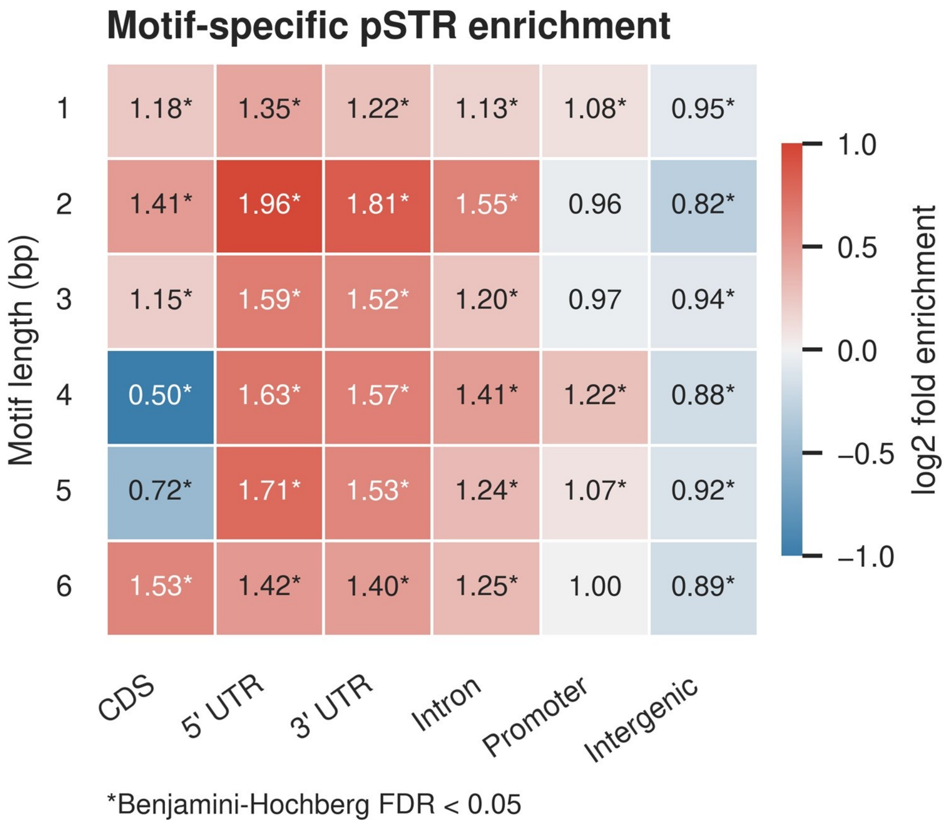


**Figure S2**: **Motif-specific enrichment of polymorphic STRs across genomic contexts.** For each motif length, cells compare the fraction of reference STR loci that are polymorphic in a genomic context with the polymorphic fraction across all genomic contexts. Cell values are fold enrichments; colors show their $\log_2$ values, with red denoting enrichment and blue denoting depletion. Asterisks indicate Benjamini–Hochberg FDR $< 0.05$ across 36 two-sided Fisher exact tests comparing each context with the other contexts of the same motif length. Promoters are strand-aware 2-kb regions upstream of annotated transcription start sites.

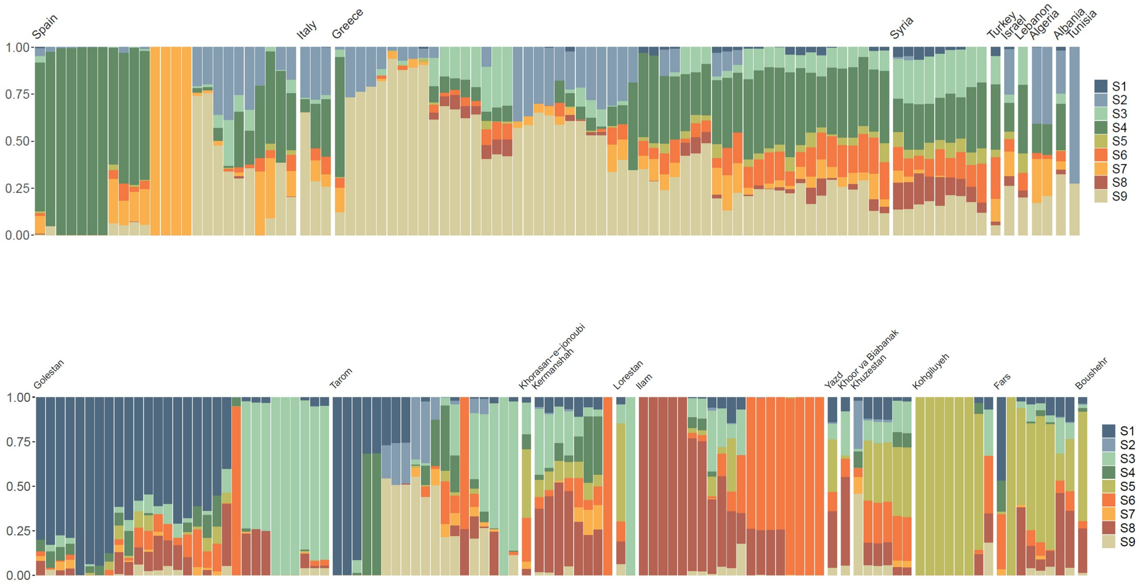


**Figure S3**: **STR-based ancestry profiles across non-Iranian and Iranian accessions.** dNMF ancestry proportions at the selected rank $K = 9$. Non-Iranian accessions (top) are grouped by country and Iranian accessions (bottom) by province. Each vertical bar represents one accession, and colors denote the nine STR ancestry components.

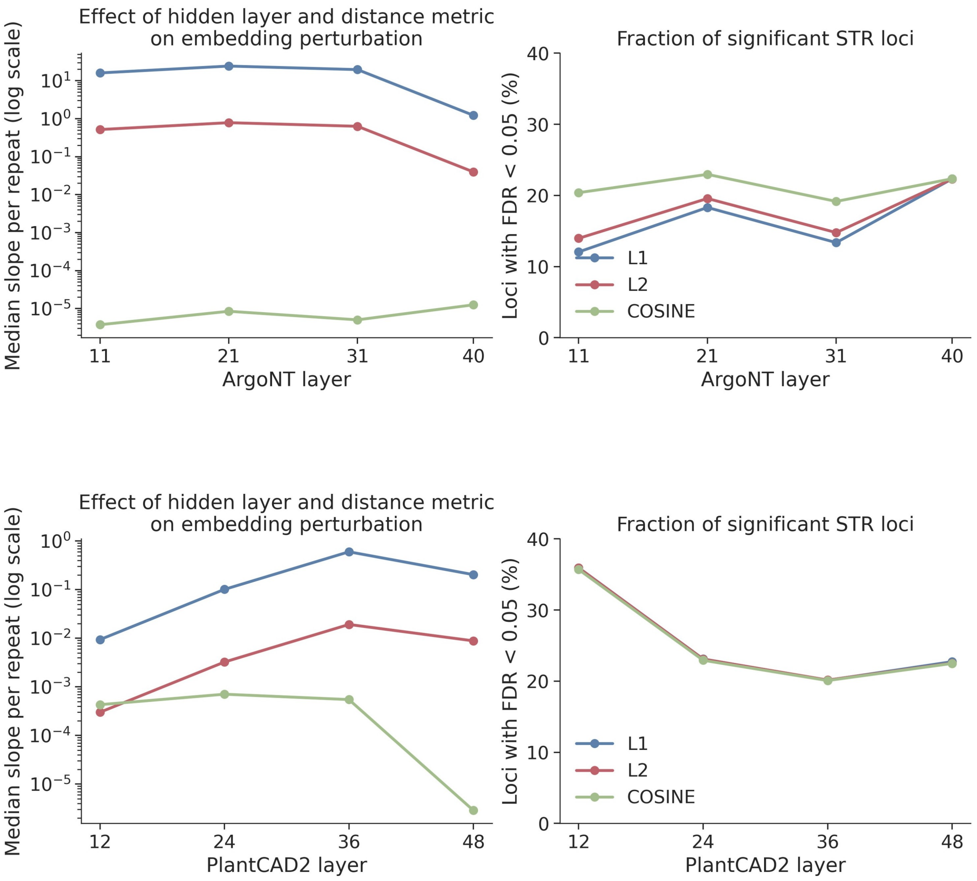


**Figure S4**: **Sensitivity of embedding perturbations to hidden layer and distance metric.** Results are shown for AgroNT (top; layers 11, 21, 31, and 40) and PlantCAD2 (bottom; layers 12, 24, 36, and 48). Left panels show the median embedding-distance slope per repeat across tested STR loci for L1, L2, and cosine distance; the vertical axes are logarithmic. Right panels show the fraction of loci significant at permutation FDR $< 0.05$. Absolute slopes are metric- and model-scale dependent. Layer 21 with whole-sequence mean L2 distance was selected for the primary AgroNT analysis, and layer 36 with the same distance definition was selected for PlantCAD2.

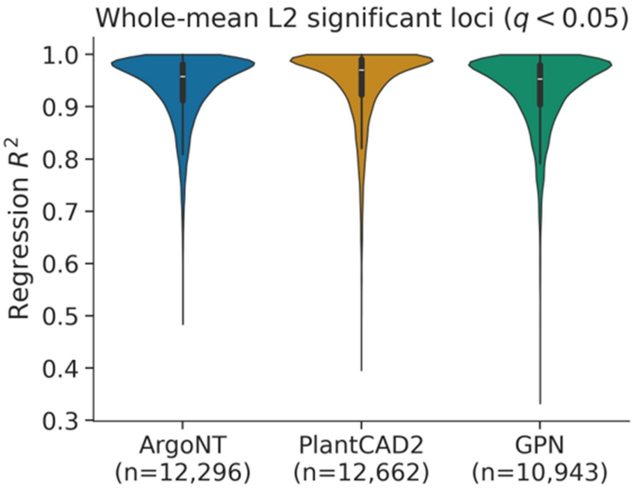


**Figure S5**: **Regression fit among embedding-sensitive STR loci.** Distribution of $R^2$ for reference-anchored whole-sequence mean L2 regressions passing permutation FDR < 0.05 in AgroNT layer 21, PlantCAD2 layer 36, and the final GPN hidden layer. Violin interiors show boxplots; sample sizes below the model names are the model-specific numbers of significant loci.

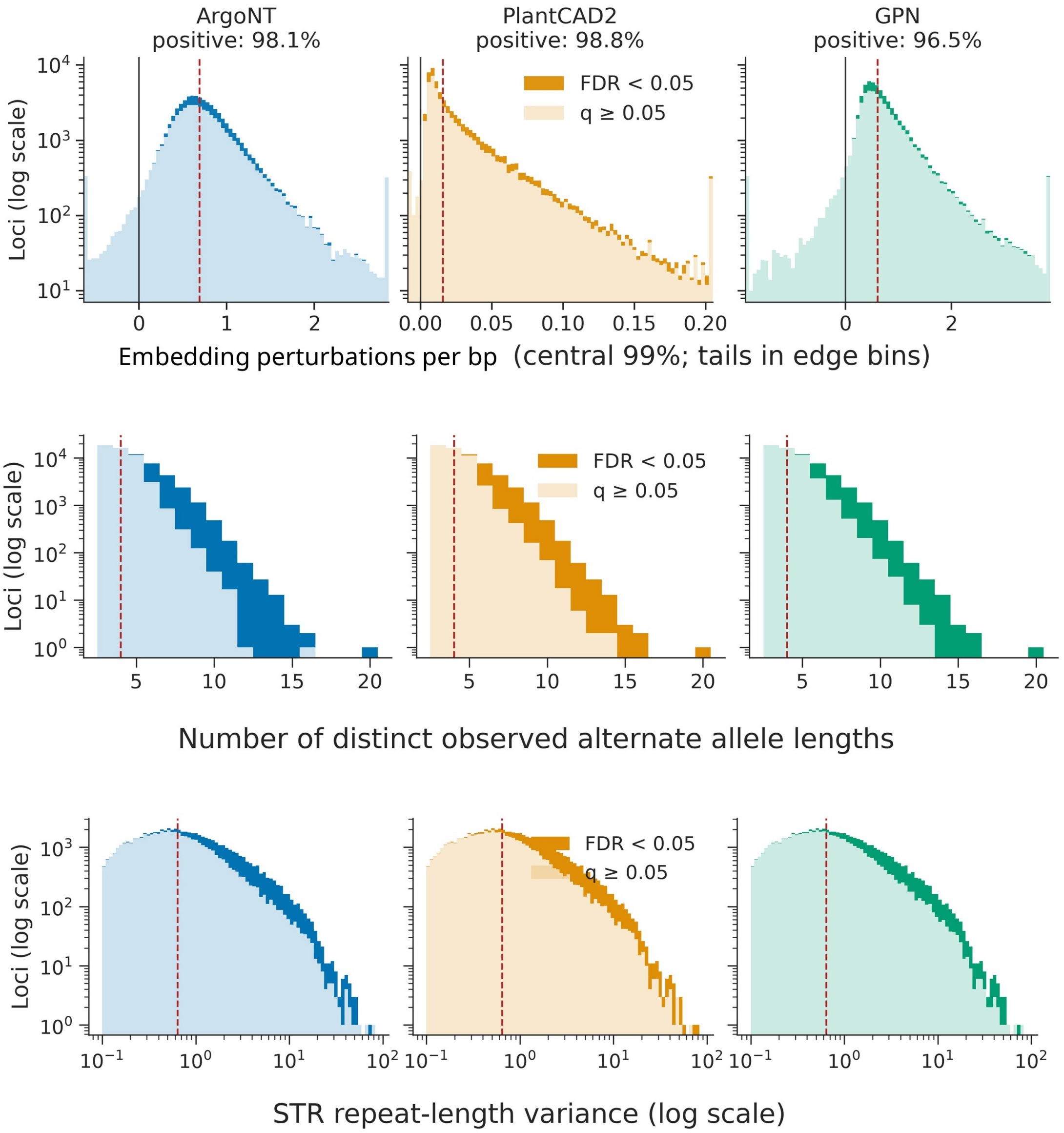


**Figure S6**: **Embedding slopes and locus properties associated with detection power.** Columns show AgroNT layer 21, PlantCAD2 layer 36, and the final GPN hidden layer for the same 62,847 tested loci. Top panels show distributions of whole-sequence mean L2 slopes per base pair, with the central 99% displayed and tails collected in edge bins; black lines mark zero and red dashed lines mark medians. Middle and bottom panels show the number of distinct observed nonreference allele lengths and population STR repeat-length variance, respectively. Dark shading denotes loci significant at permutation FDR < 0.05, and pale shading denotes nonsignificant loci.

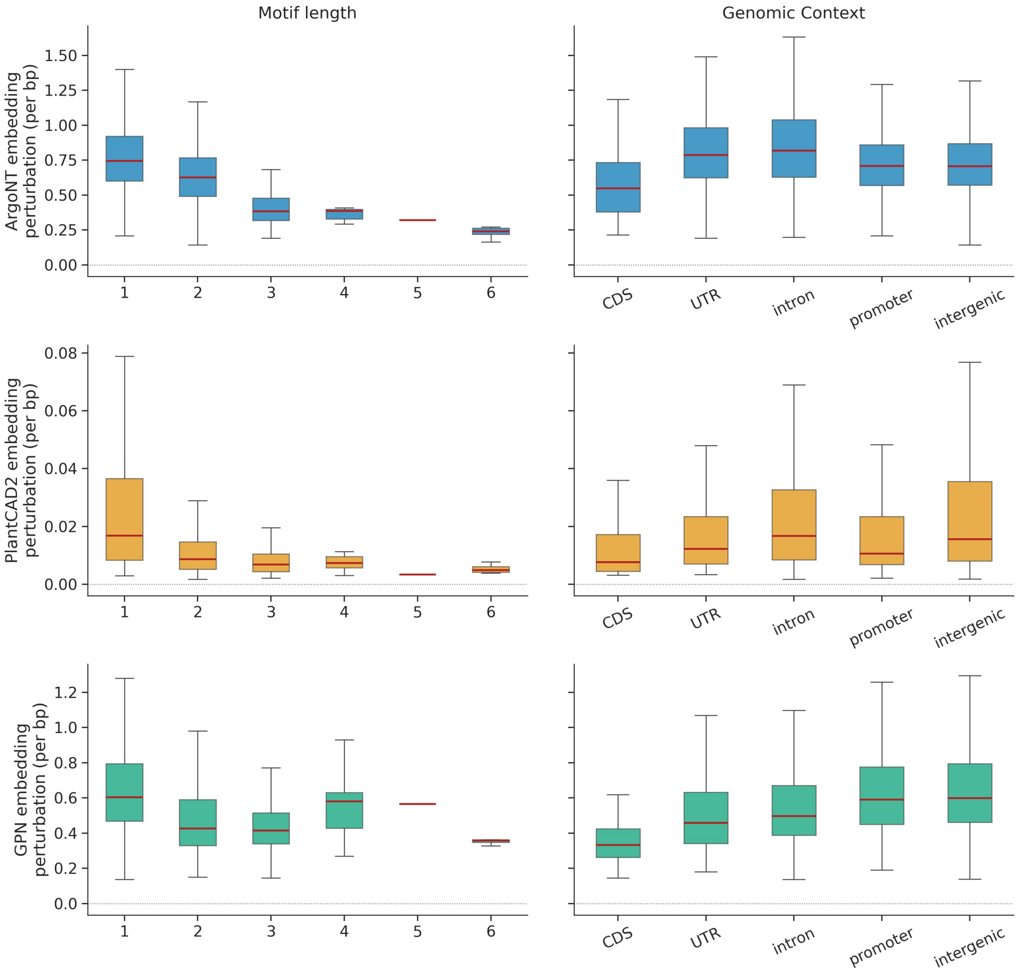


**Figure S7**: **Per-base embedding perturbation by motif length and genomic context.** Whole-sequence mean L2 slopes per base pair among loci significant at permutation FDR < 0.05 are summarized for AgroNT layer 21 (top), PlantCAD2 layer 36 (middle), and the final GPN hidden layer (bottom). Left panels stratify loci by motif length and right panels by mutually exclusive genomic context; 5′ and 3′ UTRs are combined. Boxes show interquartile ranges with median lines, whiskers extend to the most extreme observations within 1.5 interquartile ranges of the box, and more extreme observations are not displayed.

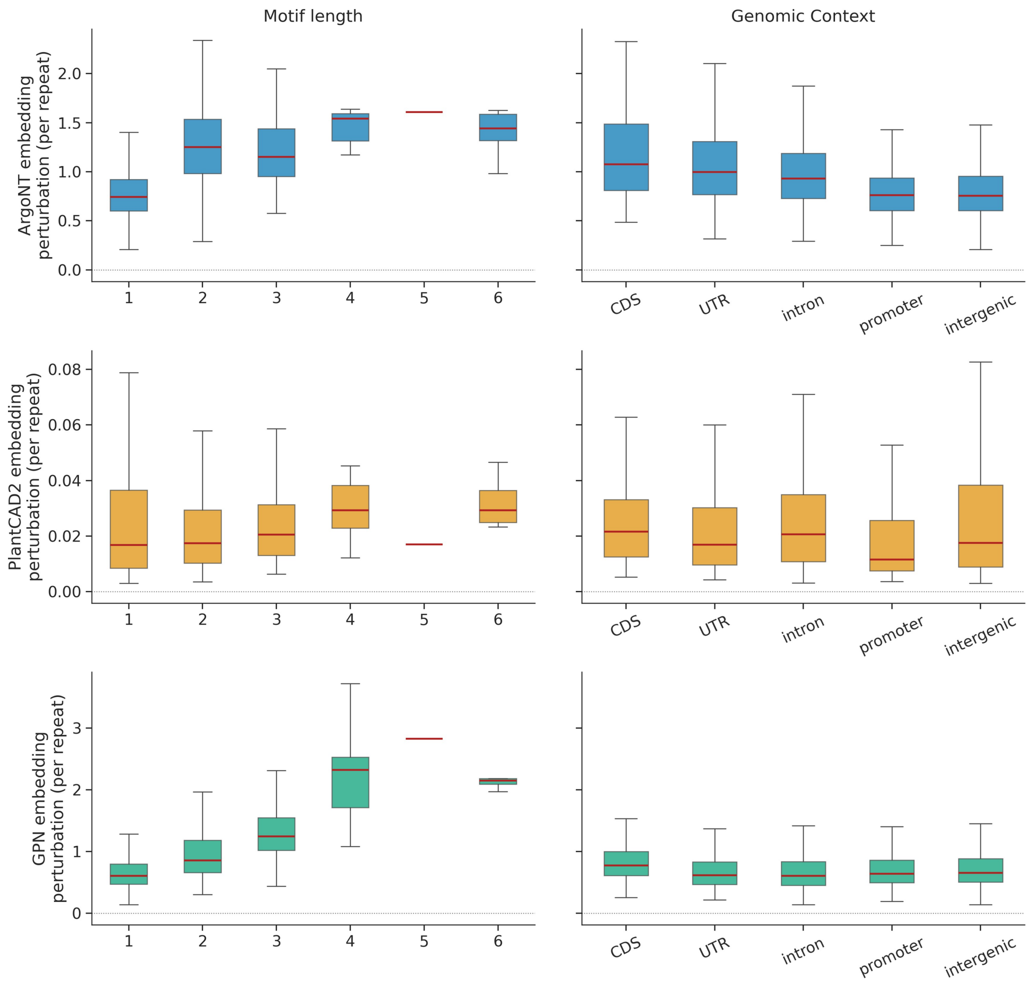


**Figure S8**: **Per-repeat embedding perturbation by motif length and genomic context.** Whole-sequence mean L2 slopes per repeat among loci significant at permutation FDR $< 0.05$ are summarized for AgroNT layer 21 (top), PlantCAD2 layer 36 (middle), and the final GPN hidden layer (bottom). Left panels stratify loci by motif length and right panels by mutually exclusive genomic context; 5′ and 3′ UTRs are combined. Boxes show interquartile ranges with median lines, whiskers extend to 1.5 interquartile ranges, and more extreme observations are not displayed. Because one repeat contributes different numbers of bases across motif classes, these distributions complement the per-base comparison in Fig. S7.

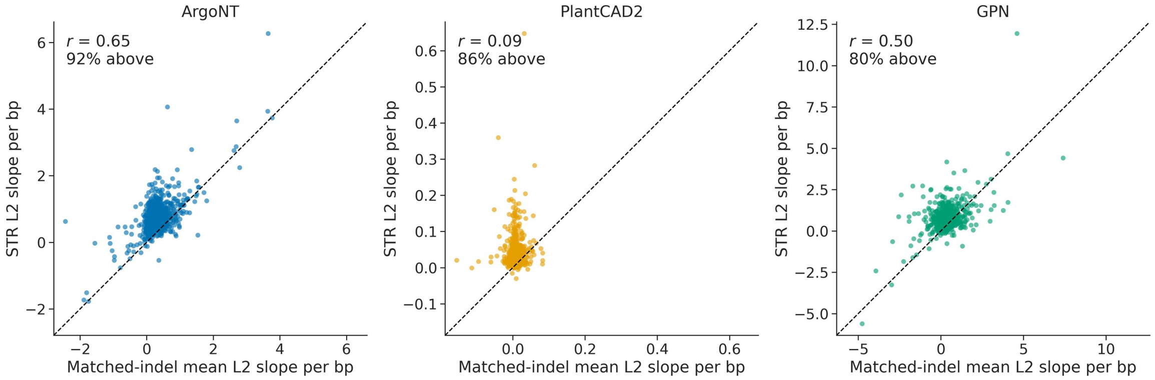


**Figure S9**: **STR embedding responses exceed length-matched nonrepetitive indel controls.** Per-base whole-sequence mean L2 slopes for 1,000 genuine STR loci are plotted against the mean slope from three deterministic nonrepetitive indel series with identical allele lengths. Panels show AgroNT layer 21, PlantCAD2 layer 36, and the final GPN hidden layer. Dashed diagonal lines indicate equal STR and control slopes; annotations report the Pearson correlation and the percentage of loci for which the STR slope exceeds the matched-control mean.

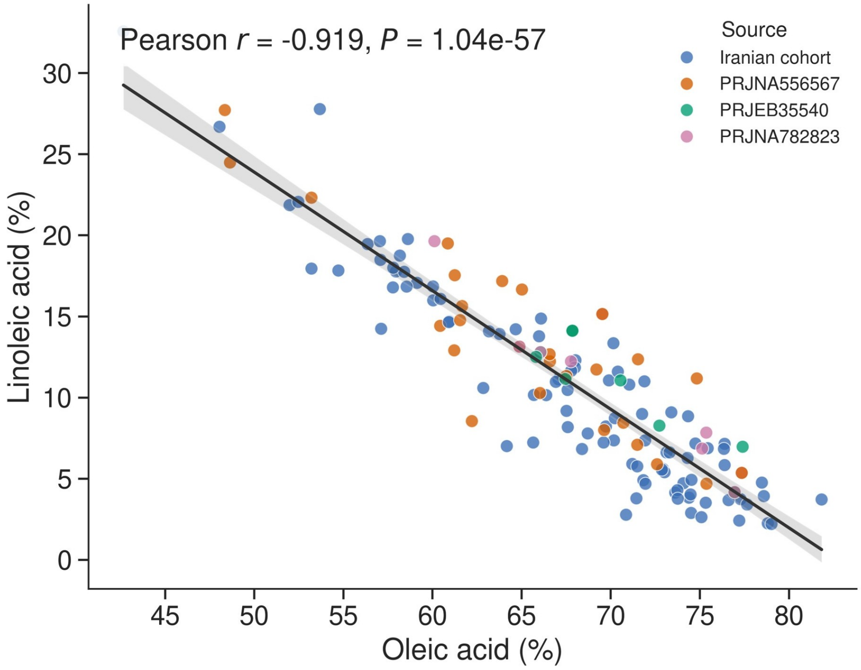


**Figure S10**: **Inverse relationship between oleic and linoleic acid content.** Oleic- and linoleic-acid measurements for 140 accessions. Colors distinguish the Iranian cohort and accessions from BioProjects PRJNA556567, PRJEB35540, and PRJNA782823. The line and shaded band show the pooled ordinary least-squares fit and its 95% confidence interval. The pooled Pearson correlation coefficient and two-sided $P$ value are reported in the panel.

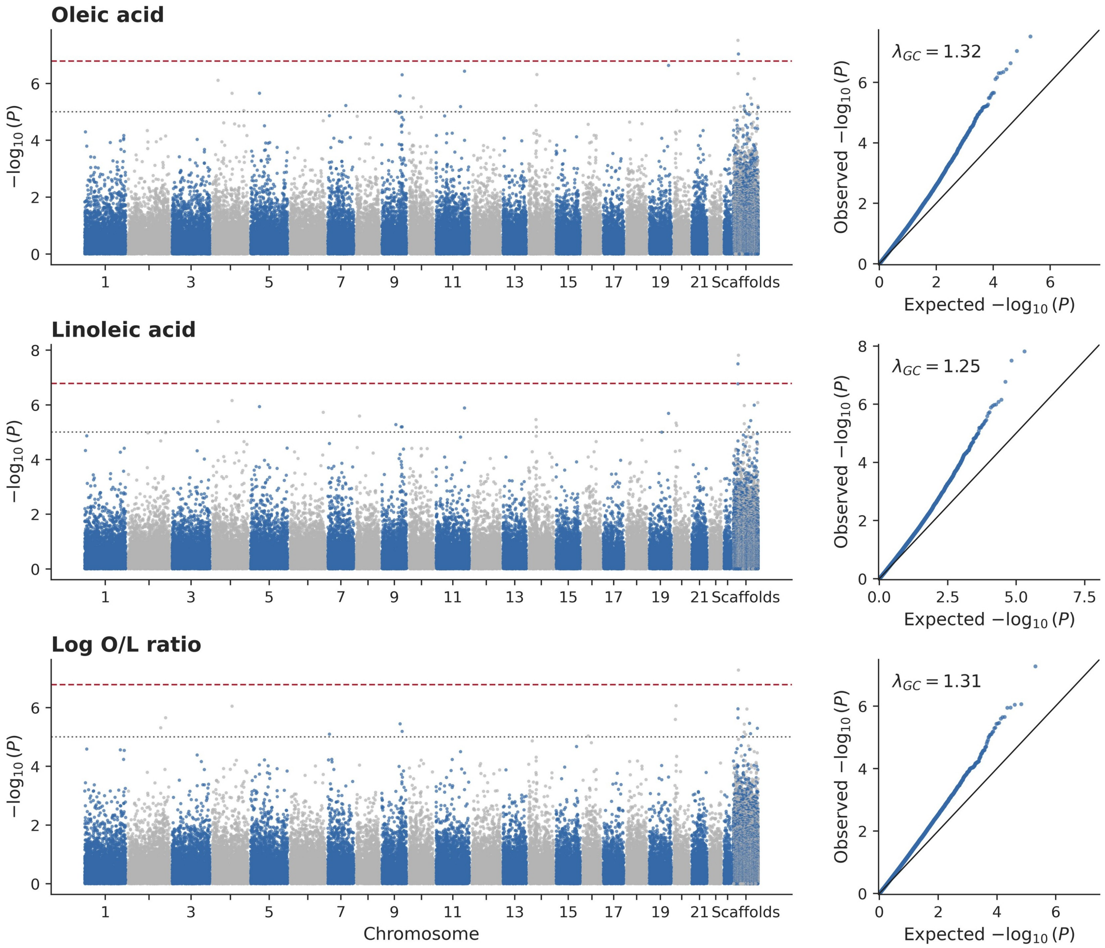


**Figure S11**: **Genome-wide STR associations with olive-oil composition.** Manhattan plots (left) and quantile–quantile plots (right) for oleic acid, linoleic acid, and the natural logarithm of the oleic/linoleic ratio. Alternating colors distinguish successive chromosomes and unplaced scaffolds. Gray dotted horizontal lines mark the suggestive threshold ($1/101{,}184 = 9.88 \times 10^{-6}$), and red dashed lines mark the study-wide Bonferroni threshold across three phenotypes [$0.05/(101{,}184 \times 3) = 1.65 \times 10^{-7}$]. Quantile–quantile panels report genomic inflation factors, $\lambda_{GC}$.

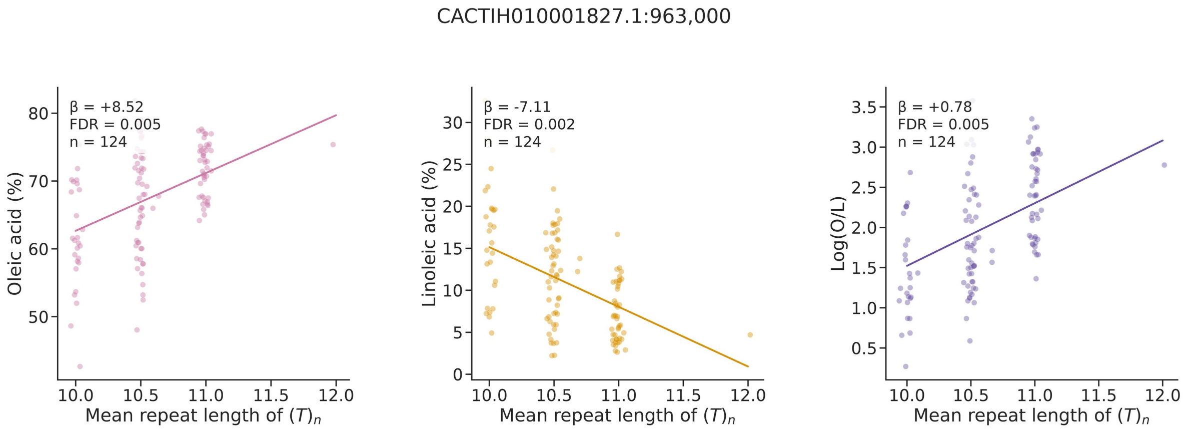


**Figure S12**: **STR locus associated with all three oil-quality phenotypes.** Associations at `CACTIH010001827.1:963,000` are shown for oleic acid (left), linoleic acid (middle), and the natural logarithm of the oleic-to-linoleic ratio, Log(O/L) (right). All three unadjusted association *P* values pass the study-wide Bonferroni threshold of $0.05/(101{,}184\times3) = 1.65\times10^{-7}$. Points are raw sample measurements with horizontal jitter. Lines have the pooled covariate-adjusted linear mixed-model slope and are centered at the raw phenotype mean. $\beta$ is the effect per one-repeat increase in mean repeat length; for oleic and linoleic acid it is measured in percentage points, whereas for Log(O/L) it is measured on the natural-log scale. Displayed FDR values are Benjamini–Hochberg adjusted separately within each phenotype.

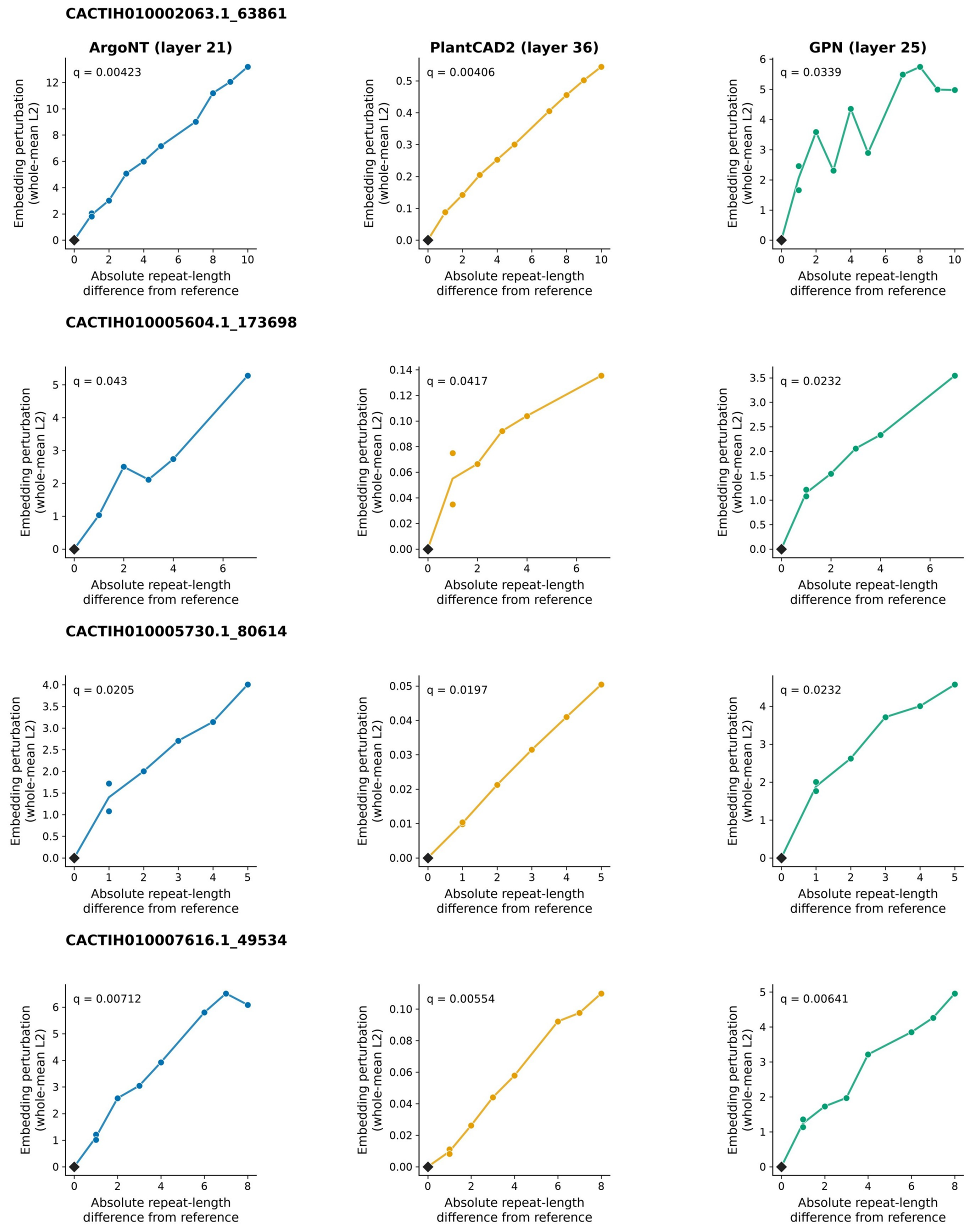


**Figure S13**: **Allele-level embedding responses at four phenotype-associated loci supported by all three models.** Rows correspond to the four prioritized STR loci and columns to AgroNT layer 21, PlantCAD2 layer 36, and the final GPN hidden layer. Whole-sequence mean L2 distance from the reference allele is plotted against absolute repeat-length difference from that reference. Black diamonds identify reference alleles, whose distance is zero by definition and which were excluded from slope estimation and permutation testing. Colored points are observed nonreference alleles, and connecting lines are visual guides only. Displayed $q$ values are model-specific Benjamini–Hochberg-adjusted permutation values.
